# Gut microbiota promotes enteroendocrine cell maturation and mitochondrial function

**DOI:** 10.1101/2023.09.27.558332

**Authors:** Alfahdah Alsudayri, Shane Perelman, Annika Chura, Melissa Brewer, Madelyn McDevitt, Catherine Drerup, Lihua Ye

**Affiliations:** Department of Neuroscience, the Ohio State University Wexner Medical Center; Department of Integrative Biology, University of Wisconsin-Madison

## Abstract

The enteroendocrine cells (EECs) in the intestine are crucial for sensing ingested nutrients and regulating feeding behavior. The means by which gut microbiota regulates the nutrient-sensing EEC activity is unclear. Our transcriptomic analysis of the EECs from germ-free (GF) and conventionalized (CV) zebrafish revealed that commensal microbiota colonization significantly increased the expression of many genes that are associated with mitochondrial function. Using in vivo imaging and 3D automated cell tracking approach, we developed new methods to image and analyze the EECs’ cytoplasmic and mitochondrial calcium activity at cellular resolution in live zebrafish. Our data revealed that during the development, shortly after gut microbiota colonization, EECs briefly increased cytoplasm and mitochondrial Ca^2+^, a phenomenon we referred to as “EEC awakening”. Following the EEC awakening, cytoplasmic Ca^2+^ levels but not mitochondrial Ca^2+^ level in the EECs decreased, resulting in a consistent increase in the mitochondrial-to-cytoplasmic Ca^2+^ ratio. The increased mitochondrial-to-cytoplasmic Ca^2+^ ratio is associated with the EEC maturation process. In immature EECs, we further discovered that their mitochondria are evenly distributed in the cytoplasm. When EECs mature, their mitochondria are highly localized in the basal lateral membrane where EEC vesicle secretion occurs. Furthermore, CV EECs, but not GF EECs, exhibit spontaneous low-amplitude calcium fluctuation. The mitochondrial-to-cytoplasm Ca^2+^ ratio is significantly higher in CV EECs. When stimulating the CV zebrafish with nutrients like fatty acids, nutrient stimulants increase cytoplasmic Ca^2+^ in a subset of EECs and promote a sustained mitochondrial Ca^2+^ increase. However, the nutrient induced EEC mitochondrial activation is nearly abolished in GF zebrafish. Together, our study reveals that commensal microbiota are critical in supporting EEC mitochondrial function and maturation. Selectively manipulating gut microbial signals to alter EEC mitochondrial function will provide new opportunities to change gut-brain nutrient sensing efficiency and feeding behavior.

## Introduction

Feeding behavior is conserved among all organisms. During development, the fetus receives its nutrient supply from its mother. Immediately after birth, the maternal nutrient supply is cut off and the infant needs to initiate the feeding process to obtain nutrients. After eating, the ingested nutrients need to be sensed, and such nutrient information will be transmitted to the rest of the body to coordinate its metabolic function [1]. Within the intestinal epithelium, a group of specialized sensory cells called enteroendocrine cells (EECs) sense the ingested nutrient information and secrete hormone molecules to regulate physiological homeostasis [2]. The EECs are critical nutrient sensors in the intestinal epithelium. They are dispersed along the digestive tract and comprised of less than 1% of the intestinal epithelium cells (IECs). However, collectively, the EECs form the largest endocrine organ in the body [2]. Most of the previous studies assessing EECs are focused on adults. It is well-known that ingested nutrients, such as fatty acids or glucose, directly stimulate the EECs by triggering a cascade of membrane depolarization, action potential firing, voltage-dependent calcium entry, and hormone-containing vesicle release [2]. Many of these EEC-secreting hormones, like Cholecystokinin (CCK) or glucagon-like peptide 1 (GLP-1), are critical in regulating satiation response and metabolism [1, 2]. In addition to the classic hormone secretion function, recent research also demonstrated that the EECs form a basal membrane process called “neuropod” that directly synapses with the vagal sensory neurons [3, 4]. Through the EEC-vagal neuronal pathway, ingested nutrient information in the gut lumen can be transmitted to the brain [4]. Further studies demonstrated that this nutrient-sensing EEC-vagal pathway is essential to drive animal’s food preference toward sugar and fat [5–7]. It is well known that the intestinal epithelium cells undergo significant remodeling during the postnatal period to adapt to the need for nutrient absorption and sensation [8]. Despite the importance of EECs in nutrient monitoring, gut-brain nutrient sensing, feeding behavior, and systemic metabolic regulation, little is known about how environmental factors regulate EEC maturation and function during the postnatal developmental period.

Following birth, newborn babies are rapidly colonized by microbial organisms [9]. These microbial organisms start to assemble the functional microbial community that plays important roles in the development of the infant [9]. Previous studies revealed that microbiota colonization during the early postnatal phase is critical in promoting intestinal epithelium maturation and remodeling [10–12]. Numerous pieces of evidence also suggest that gut microbiota are important in regulating nutrient absorption, metabolism, and infant growth [13–15]. Research from animal models and clinical studies suggest that gut microbiota are critical in modulating feeding behavior, including appetite and food choice [16]. However, little is known about how gut microbiota interacts with EECs and regulates the EECs’ function. Whether and how gut microbiota interacts with EECs to modulate postnatal physiological homeostasis remains unknown.

Mitochondria is the essential organelle that provides ATP to sustain cellular function. The mitochondria emerge as a key player in coordinating cellular metabolism, cell differentiation, and regulating intestinal epithelium homeostasis [17–19]. In the nervous system, mitochondria are important in supporting neuronal vesicle secretion, neuronal synaptic transmission, and neuronal network formation [20]. Research in pancreatic endocrine cells also revealed that mitochondrial function is essential for pancreatic beta-cell hormonal secretion [21]. Interestingly, the pancreatic islet’s mitochondrial activity is inhibited in diabetic conditions [21, 22]. Little is known about the physiological roles of the mitochondrial function in EECs, and how environmental factors regulate EEC mitochondrial activity *in vivo*.

A major challenge in studying how environmental factors, such as gut microbiota, regulate EEC physiology has been the lack of in vivo techniques to study this rare intestinal epithelium population in the intact animal setting. Historically, these cells have been studied by measuring the circulating EEC-secreting hormones [2]. However, many EEC hormones have very short half-lives and the plasma hormone level does not mirror the EEC function nor the real-time EEC activity [23]. EEC activity has also been studied via cell culture or organoid culture systems. However, a cell or organoid culture is not able to mimic the dynamic and complex intestinal luminal factors that shape the EECs. It is also difficult to study how EECs communicate with neighboring cells or distant organs, like the brain, using the in vitro culture system.

In this study, we utilized the zebrafish model to examine how commensal microbiota affect EEC maturation and function. Using an innovative approach to direct images and track the EEC cellular and mitochondrial calcium activity in live zebrafish during development, our results revealed that the EEC morphology, cellular, and mitochondrial activity is dynamically regulated during the EEC maturation process. Importantly, our results revealed that gut microbiota play critical roles in promoting EEC maturation and mitochondrial function.

## Results

### Gut microbiota alters EEC subtypes

Previous studies, including ours, demonstrated that similar to mammals, the zebrafish EECs have diverse subtypes [24, 25]. A recent zebrafish intestine epithelium single-cell RNA sequencing dataset further revealed the five EEC subtypes in the zebrafish larvae characterized by their distinct hormone expression profiles (Fig. S1A-B) [26]. EEC1 is characterized by the expression of the hormonal genes: pancreatic polypeptide b (*pyyb*), somatostatin 2 (*sst2*), and ghrelin (*ghrl*) (Fig. S1A-B). EEC2 expresses the hormonal genes: preproglucagaon a (*gcga*), the gene that encodes Glucagon (Gcg) or Glucan-like-pepetide 1 (GLP-1), vasoactive intestinal polypeptide b (*vipb*), and insulin-like 5 (*insl5*) (Fig. S1A-B). EEC3 expresses the hormonal genes: calcitonin related polypeptide alpha (*calca*) and neuromedin b (*nmb*) (Fig. S1A-B). EEC4 expresses the hormonal genes: cholecystokinin a (*ccka*) (Fig. S1A-B). EEC5 uniquely expresses the following hormonal genes: brain-derived neurotrophic factor (*bdnf*), adenylate cyclase-activating peptide-1 a (*adcyap1a*), preproenkephalin a (*penka*), and tryptophan hydroxylase 1 b (*tph1b*), the enzyme that synthesizes serotonin (Fig. S1A-B). The EEC5 also highly and uniquely expresses the Transient receptor potential ankyrin 1 b (Trpa1b). These Trpa1+EECs that are characterized in our previous studies sense microbial stimulants and are critical in regulating gut motility and intestinal homeostasis [27]. Some of these EEC markers can be labeled via immunofluorescence staining and transgenic approaches (Fig.S1C-K). Our results confirmed the hormonal expression profiles in different EEC subtypes that are revealed in the single-cell RNA sequencing above (Fig. S1C-K). Moreover, consistent with previous studies, our data revealed that the distribution of the EEC subtypes exhibits regional specificity (Fig. S1C-K). For example, the PYY+EECs are exclusively in the proximal intestine, while the Trpa1+EECs are distributed along the whole digestive tract. Interestingly, the Trpa1+EECs (EEC5) appeared to have heterogeneity. As the proximal intestinal Trpa1+EECs express both Enk and Serotonin (Fig. S1C-K). However, the middle and distal intestinal Trpa1+EECs do not express ENK but only express serotonin (Fig. S1C-K).

Next, we investigated how commensal gut microbiota affect EEC subtype specification via the zebrafish gnotobiotic approach (Fig. 1A) [28]. The zebrafish were derived as germ-free at 0 day post fertilization (dpf) [28]. At 3dpf, the zebrafish are maintained as germ-free (GF) or colonized with commensal microbiota (Conventionalized, CV). The gnotobiotic zebrafish were fed from 3dpf to 7dpf, and the zebrafish were fixed at 7dpf for immunofluorescence staining. Our results revealed that commensal microbiota colonization did not significantly alter the percentage of PYY+ EECs and ENK+EECs in the proximal intestine (Fig. 1B-E’, J-K). However, commensal microbiota colonization decreased the number of Gcg/GLP-1+EECs in the proximal intestine and Trpa1+EECs in the distal intestine (Fig. 1F-I, L-N).

**Figure 1.**
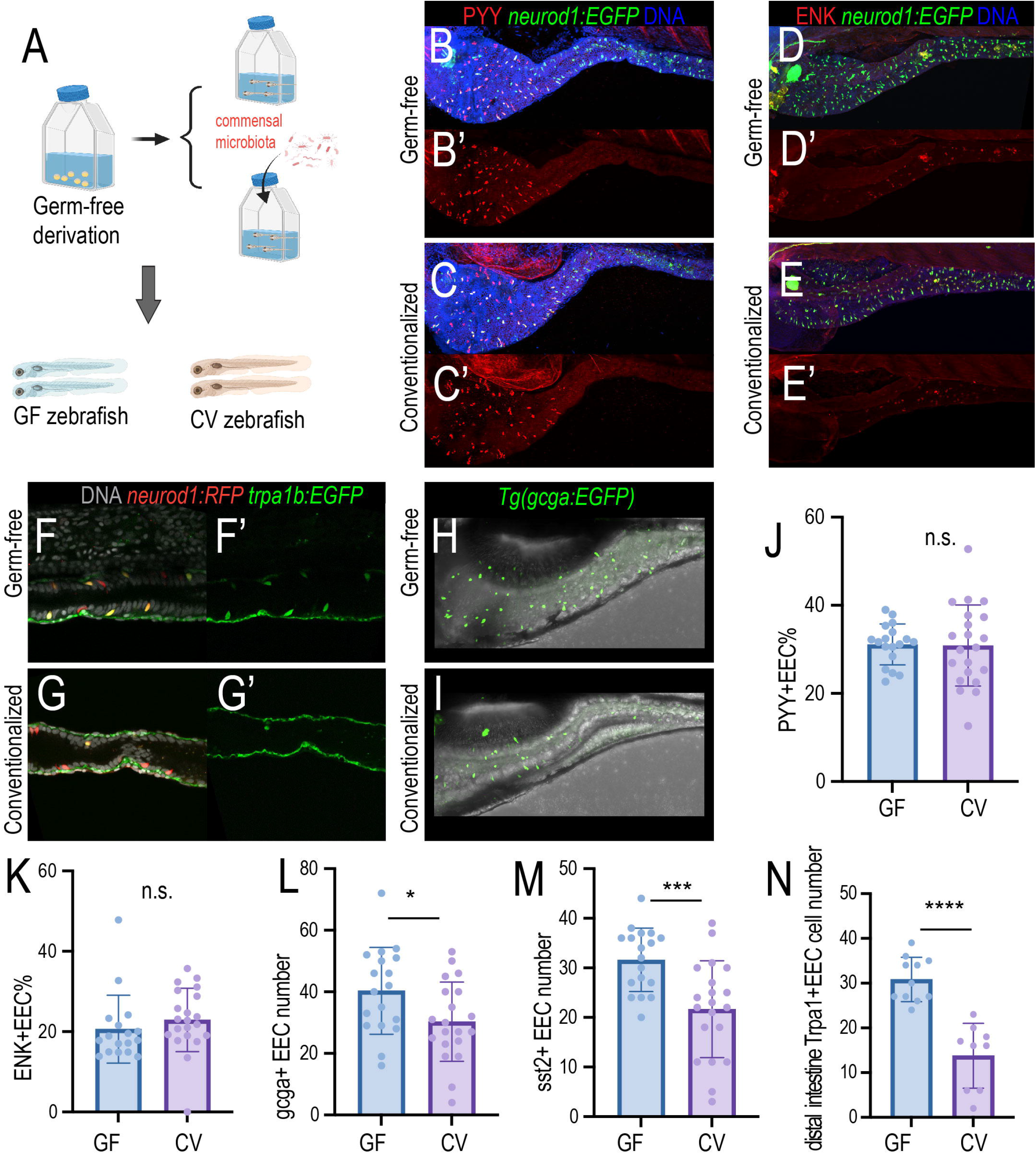
Gut microbiota modulates the EEC subtype. (A) Gnotobiotic zebrafish experimental procedure to examine the effects of gut microbiota on EEC subtype formation. Commensal microbiota was colonized at 3pdf and the zebrafish were fixed at 7dpf for immunofluorescence staining. (B-C’) Confocal projection of the representative germ-free (GF) and conventionalized (CV) zebrafish intestine. The total EECs were labeled by the *Tg(neurod1:EGFP)* transgene (green), and the PYY+EECs were labeled via the PYY antibody. (D-E’) Confocal projection of the representative germ-free (GF) and conventionalized (CV) zebrafish intestine showing the ENK+ EECs. (F-G’) Confocal projection of the representative germ-free (GF) and conventionalized (CV) zebrafish intestine showing the Trpa1+ EECs in the distal intestine. (H-I) Confocal projection of the representative germ-free (GF) and conventionalized (CV) zebrafish intestine showing the gcga+ EECs. (J-N) Quantification of the PYY+EECs, ENK+EECs, gcga+EECs, sst2+EECs and the Trpa1+EECs in GF and CV zebrafish. Student t-tests were used for statistical analysis. Each dot represents individual zebrafish. *p<0.05, *** p<0.001, **** p<0.0001.

### Gut microbiota promotes EEC maturation and mitochondrial function

To further understand how commensal microbiota modulate EECs in zebrafish, we used Flow Activated Cell Sorting (FACS) to isolate EECs from GF and CV zebrafish and performed transcriptomic analysis (Fig. 2A). For each gene, the fold change in response to gut microbial status (CV vs GF) and fold change in response to the cell fate (EEC vs other IEC) was plotted (Fig. 2B). Our results demonstrated that there is a weak but significant positive correlation between genes that are enriched in EECs and the genes that are upregulated in the CV condition(Fig. 2B). Within the genes that are significantly upregulated in CV, about 74.5% of them are enriched in the EECs (Fig. 2C). We then plotted the conserved EECs’ signature genes that are shared among zebrafish, mice, and humans [27] (Table 1). Our results indicate that about 72% of those conserved EEC signature genes are upregulated in CV (Fig. 2D, Table 1). Within those conserved EEC genes, many of them are associated with EEC cell membrane potential regulation and vesicle secretion (Fig. 2E). The CV condition also significantly upregulates the chromogranin A gene (*chga*), which labels the mature EECs (Fig. 2E)[29]. Therefore, consistent with the immunofluorescence results above, the transcriptomic analysis indicates that gut microbiota promote EEC cell fate and maturation. Next, we performed Go Term analysis of the genes that are significantly upregulated in CV and the genes that are significantly upregulated in GF using the Metascape gene analysis tool (Fig. 2F-G). Interestingly, in the genes that are significantly upregulated in GF EECs, top Go Term includes gene functions related to adhesion, migration, and actin filament-based processes (Fig. 2G). Within the CV upregulated genes, several Go Terms associated with mitochondrial function are enriched (Fig. 2G).

**Figure 2.**
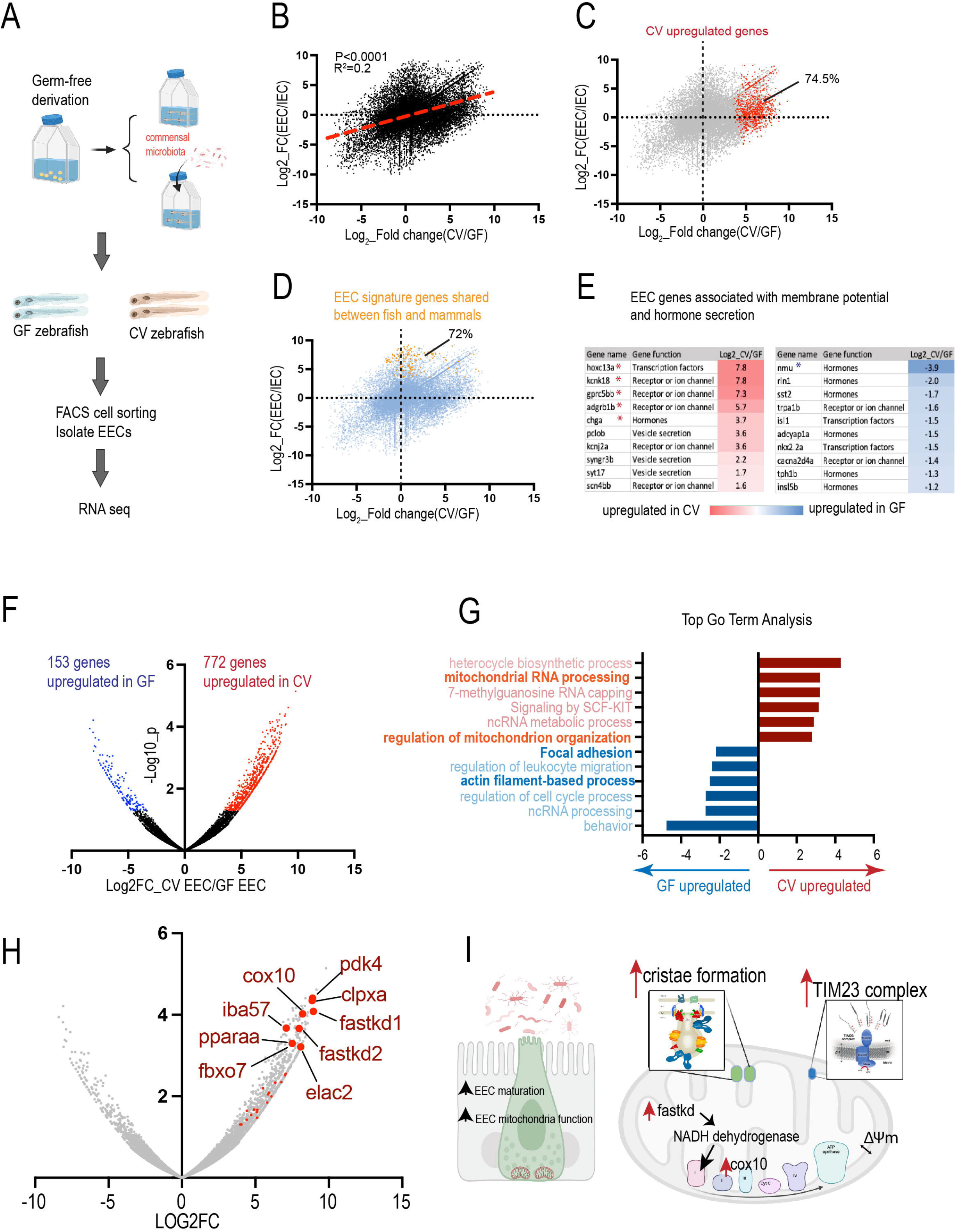
Gut microbiota promotes EEC maturation and mitochondrial function. (A) Transcriptomic analysis of the FACS sorted EECs from GF and CV zebrafish. (B) Positive correlation between the genes that are upregulated in CV (X-axis) and the genes that are enriched in EECs (Y-axis). (C) Among the genes that are significantly upregulated in CV (red color), 74.5% are enriched in the EECs. (D) 72% of the EECs signature genes shared between zebrafish and mammals are upregulated in CV. (E) The differential expression of the EEC signature genes that encode hormone peptides or are involved in membrane potential in GF and CV conditions. * Indicates that the genes are significantly upregulated in the GF or CV conditions. (F) The volcano plot shows the genes that are significantly upregulated in CV or GF. (G) Go-term analysis of the CV or GF upregulated genes. (H) The volcano plot shows the genes that are involved in mitochondrial function. Many of the genes that are associated with mitochondrial regulation are among the most significantly upregulated genes in CV EECs. (I) The model figure shows that commensal microbiota colonization promotes EEC maturation and EEC mitochondrial function.

Next, we annotated all of the genes associated with different aspects of mitochondrial function (Table). We found that almost all of the genes in the FASTK mitochondrial RNA binding family, TIM23 complex, and mitochondrial contact site and cristae organizing system are upregulated in CV (Table 1). The mitochondrial DNA encodes 13 proteins that are critical for electron transport chain reactions [30]. The FAS-activated serine/threonine kinase family (FASTK) is located in the mitochondrial matrix and plays an important role in processing RNA transcribed from the mitochondrial DNA [31]. The FASTK gene family is essential for synthesizing the components of the electron transport chain [31]. Within the FASTK family, previous studies showed that FASTK and FASTKD2 increase the NADH dehydrogenase transcripts and promote mitochondrial respiration specifically [31, 32]. FASTKD2 is one of the most significant genes upregulated in CV EECs (Fig. 2H). Mitochondria acquire most of their protein from the cytosol [33]. The TIM23 complex is essential for translocating cytosolic preprotein into the mitochondrial matrix across the mitochondrial membrane [33]. Within the mitochondria, the inner membrane forms invaginations known as cristae. The cristae are very specialized structures that support respiration [34]. Our results indicate that many genes that are associated with cristate organization are upregulated in CV EECs (Table1), suggesting that commensal microbiota colonization increases EEC mitochondrial respiration function. In addition to the genetic pathways above, our results also indicate that cox10 (an important component of mitochondrial respiration for complex III) is among the most significantly upregulated CV EEC genes (Fig.2H). Together, our transcriptomic data indicates that gut microbiota promote EEC maturation and mitochondrial function by increasing different genetic pathways that are involved in mitochondrial respiration (Fig. 2I).

### Immature EECs contain active filopodia structures at the basal lateral membrane

Our RNA sequencing results revealed that GF zebrafish exhibit increased gene expression related to their actin filament. We then used a *Tg(neurod1:lifeActin-EGFP)* transgenic zebrafish model [24] to examine EEC actin filament dynamics. The 3dpf and 6dpf proximal intestines of the fixed zebrafish samples were examined. The zebrafish EECs start to form at around 60 hours post fertilization. At around 3dpf, the zebrafish hatched from their chorion. The gut lumen opens and gut microbiota starts to colonize the intestine. Previous studies demonstrated that the intestinal epithelium cells, including EECs, are highly polarized and contain a dense actin network in the microvilli at the apical brush border [24]. To our surprise, our data showed that at 3dpf, almost all of the zebrafish EECs exhibit complex actin filament protrusions at the base (Fig. 3A-A’). Interestingly, we did not detect active basal actin filament in other IECs at 3dpf zebrafish (Video 1). This indicates that the formation of the basal actin filaments is not associated with the general intestinal epithelium development process. It is a unique phenomenon that involves immature EECs. By 6dpf, the EECs’ basal actin filaments disappeared. Most EECs exhibited typical spindle-type morphology with a flat base (Fig. 3B-B’). To further examine the EEC actin filament dynamic change, we performed live imaging of the EECs at 3dpf *Tg(neurod1:lifeActin-EGFP)* in zebrafish and traced the same fish to 6dpf. Consistently, we observed that at 3dpf, almost all the EECs have complex actin filopodium filaments in the basal lateral portion, and the EEC extends and retracts their filopodium filaments constantly (Fig. 3C-D’) (video 2). However, in the same zebrafish, by 6dpf, the EECs do not have actin filaments at the base but contain a high lifeActin-EGFP signal at the apical brush border (Fig. 3E-F’) (video 3). Our data revealed for the first time that the immature EECs have an active filopodia process at the basal lateral membrane. When the EECs start to develop and mature, they become more polarized and lose their basal filopodia process. In general, filopodia are an antenna for cells to probe their environment [35]. The function of the immature EEC filopodia and the molecular mechanisms that regulate the EEC filopodia formation require further investigation.

**Figure 3.**
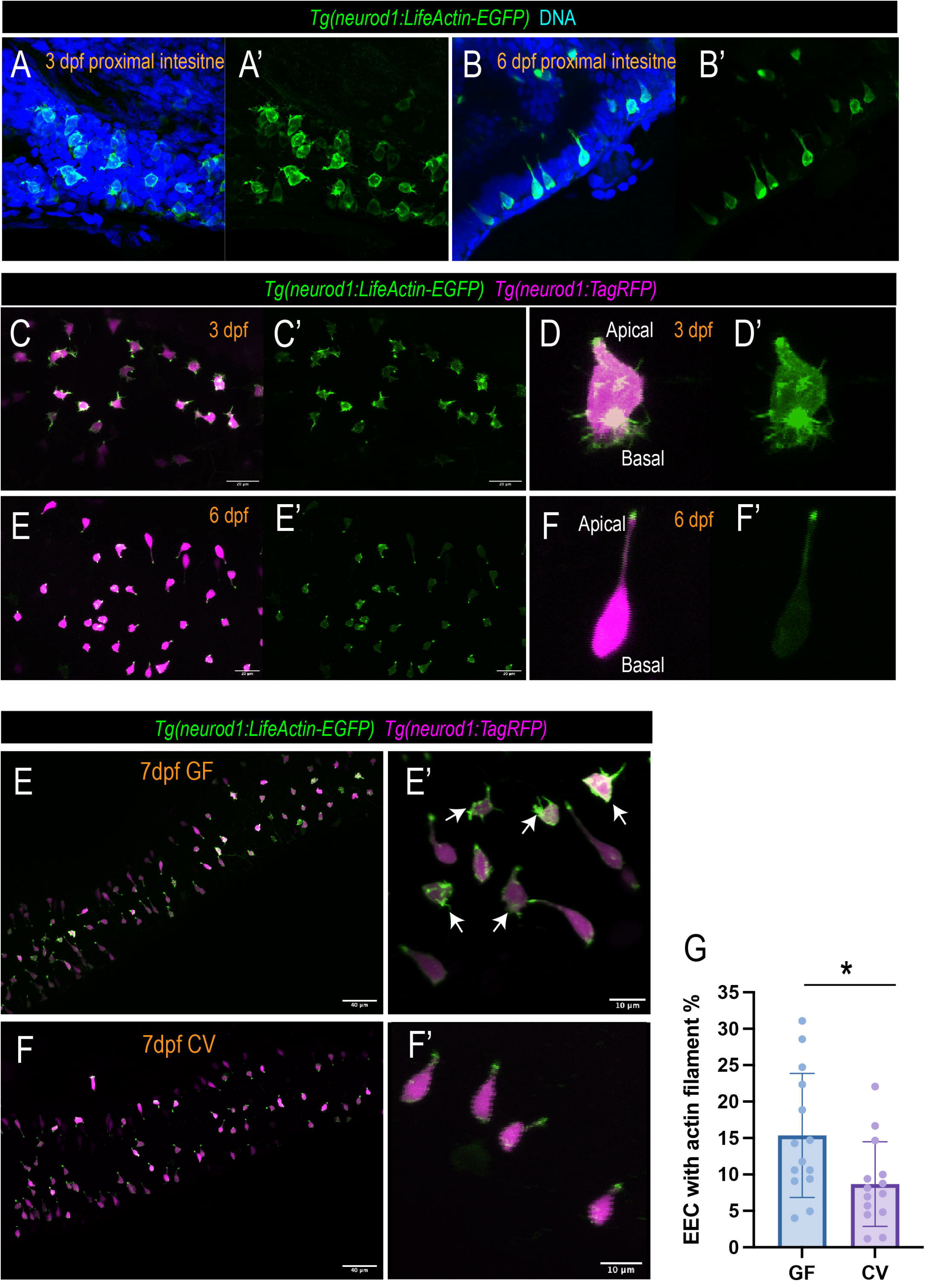
EECs change morphology during development in a microbial-dependent manner. (A-B’) Confocal projections of the proximal intestine *Tg(neurod1:lifeActin-EGFP)* zebrafish at 3dpf and 6dpf in the fixed samples. The EECs in the 3 dpf but not 6 dpf intestine exhibit thin actin filaments in the basal lateral membrane. (C-E’) Live imaging traces the same zebrafish’s EECs at 3dpf and 6dpf. The EEC actin filaments are labeled via the *Tg(neurod1:lifeActin-EGFP)*. (D-F’) Zoom out view showing the typical 3dpf EEC and 6dpf EEC. Note that at 3dpf, active actin filaments are observed at the basal lateral membrane. At 6dpf, the actin filaments are only enriched in the apical brush border. (E-F’) Confocal projection of the 7 dpf GF and CV zebrafish EECs. The EEC actin filaments are labeled via the *Tg(neurod1:lifeActin-EGFP)*. Note that the active basal lateral actin filaments remained in a subpopulation of EECs in GF zebrafish. (G) Quantification of the percentage of EECs with actin filaments in GF and CV conditions. Each dot represents an individual zebrafish. Student-T test was used in G. *P<0.05.

To investigate how gut microbiota regulate the EEC filopodia process and maturation, we generated GF and CV *Tg(neurod1:lifeActin-EGFP)* zebrafish. We imaged their proximal intestine at 7dpf. Our results demonstrated that about 15% of GF zebrafish EECs in the proximal intestine still have active filopodia actin filament at the base at 7dpf (Fig. 3E, G). The ratio of the EECs with active filopodia actin filament is significantly reduced in CV zebrafish (Fig. 3F, G). This data suggests that certain microbial cues promote EEC maturation and facilitate the disappearance of the EEC filopodia actin filament. This data is also consistent with the RNA seq results above, regarding the GF zebrafish EECs’ increased gene expression associated with the actin-filament based process (Fig. 1G).

Previous mice studies suggested that EECs form an extended membrane process called “neuropod” to connect with the nervous system [3, 36]. Interestingly, we observed that some EECs in 7dpf zebrafish intestines formed an extended membrane process in the basal membrane that morphologically resembles the mammalian neuropod of EECs (Fig. S2C-D’). The extended membrane process is distinct from the thin actin filopodia filament and is not detected in 3dpf zebrafish EECs. The neuropod-like EECs are rare in the GF zebrafish intestine (Fig. S2A-B’). The CV zebrafish exhibit a significantly higher percentage of neuropod-like EECs in the intestine (Fig. S2E). Our data suggests the formation of the neuropod-like structures in mature EECs may require certain microbial cues.

### Commensal microbiota promotes the formation of mitochondria hot spots in the EEC basal membrane

Next, we seek to determine how gut microbiota regulate the EECs’ mitochondria. We used a *Tg(neurod1:mitoEOS)* transgenic zebrafish model to visualize the EECs’ mitochondria [37]. In this model, the green fluorescent EOS protein contains a mitochondrial tag that is expressed in EECs to label their mitochondria. Using this model, we can trace the EEC mitochondrial abundance and intracellular mitochondrial distribution in live zebrafish over time. Our results revealed that, from 3dpf to 5dpf, the zebrafish EEC did not significantly increase the mitochondrial abundance (Fig. S3A). However, at 6dpf, EECs exhibit higher mitochondrial abundance compared with 3dpf-5dpf EECs (Fig. S3A). Interestingly, at 3dpf, the mitochondria are evenly distributed within the EECs, and the mitochondrial contents near the basal lateral membrane are low (Fig. 4A-B’, E). By 6dpf, most EECs exhibit hot spot mitochondrial distribution patterns (Fig. 4C-D’, E). High mitochondrial contents are found at the base of EECs, presumably at the sites where EECs secrete vesicles (Fig. 4C-D’, E). When we compare GF and CV zebrafish, commensal microbiota colonization did not increase the EEC mitochondrial abundance (Fig. S3B-C). However, the commensal microbiota colonization promotes the formation of mitochondrial hot spots at the basal membrane (Fig. 4F-H) and the CV zebrafish had higher mitochondrial contents near the basal membrane (Fig. 4F-G, I).

**Figure 4.**
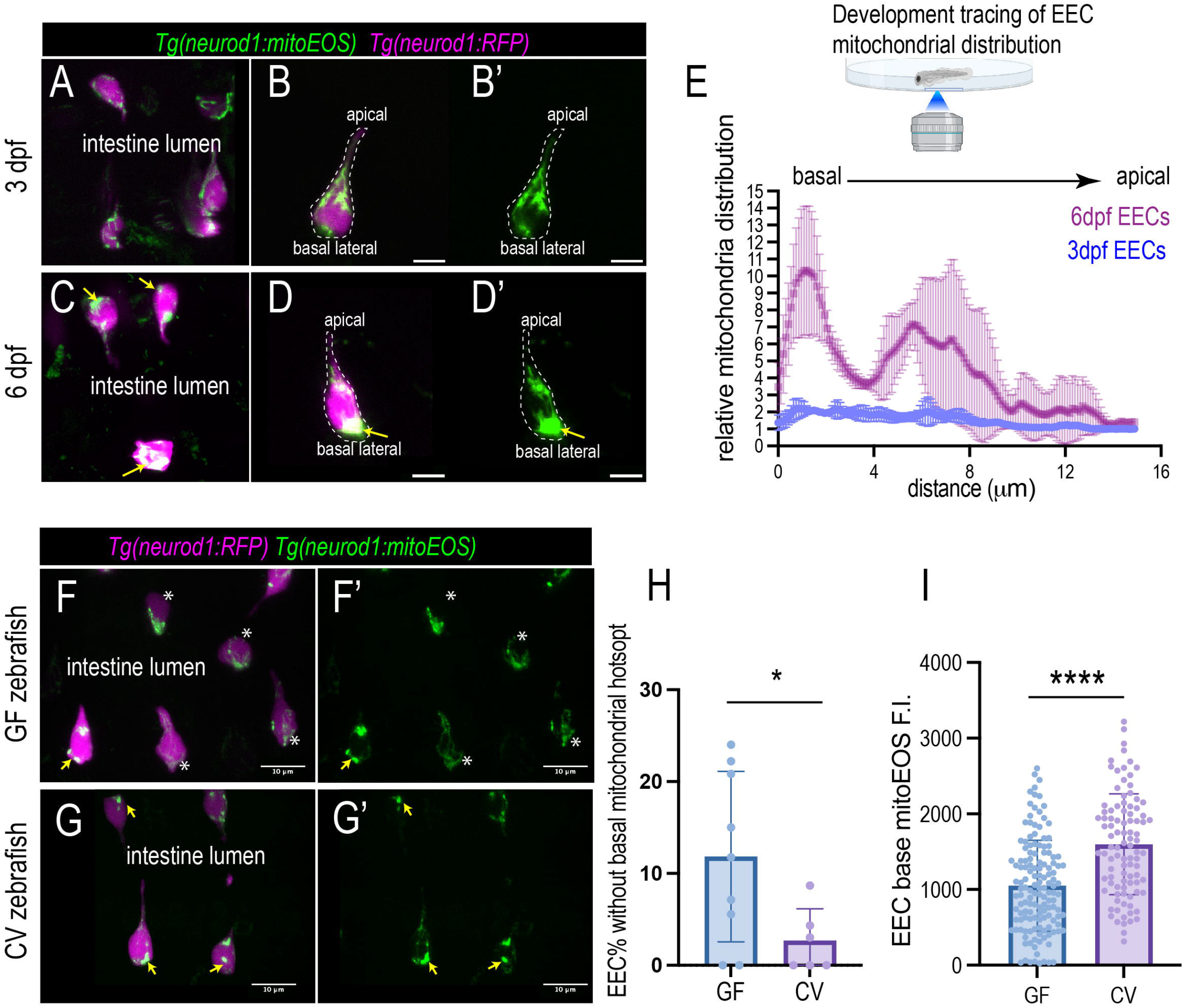
Commensal microbiota colonization promotes mitochondria accumulation at the EEC basal lateral membrane. (A-D’) In vivo imaging to trace the EEC mitochondrial abundance and the intracellular mitochondria distribution in the same zebrafish. (A-B’) Confocal projections of the typical EECs at 3dpf zebrafish proximal intestine. (C-D’) Confocal projections of the typical EECs at 6dpf zebrafish proximal intestine. The mitochondria are labeled via the *Tg(neurod1:mitoEOS),* and the EECs are labeled via the *Tg(neurod1:RFP)*. Note that at 3dpf, the mitochondria are evenly distributed in the EEC cytoplasm, and the mitochondria contents at the basal lateral membrane are low. At 6dpf, the mitochondria distribution exhibits a hot spot pattern. In most EECs, a spot at the basal lateral membrane (yellow arrows in C-D’) contains highly abundant mitochondria. (E) Quantification of the relative mitochondrial fluorescence intensity within the EECs in the same zebrafish at 3dpf and 6dpf. 5 representative EECs from the same 3dpf and 6dpf zebrafish were used to perform the data analysis. (G-H’) Confocal projections of the typical EECs in GF and CV zebrafish. Note that many EECs in GF zebrafish have low mitochondria contents at the basal lateral membrane (white stars in G and G’). Most EECs in CV zebrafish exhibit hot spot basal lateral membrane mitochondrial distribution patterns (yellow arrows in H and H’). (I) Quantification of the percentage of EECs without basal mitochondrial hotspots in 7dpf GF and CV zebrafish. Each dot represents an individual zebrafish. (I) Quantification of the mitochondrial fluorescence intensity at the basal membrane in 7dpf GF and CV zebrafish. Student T-test was used in H and I. * P<0.05, **** P<0.0001.

### Mature EECs increase mitochondrial activity

To analyze the dynamic change of EECs’ cellular and mitochondrial activity during their development and maturation process, we used the *Tg(neurod1:Gcamp6f); Tg(neurod1:mito-RGECO)* dual transgenic model [37]. In this model, the green fluorescent calcium indicator protein Gcamp6f is expressed in the EEC cytoplasm. A red fluorescent calcium indicator protein RGECO that contains a mitochondrial tag is expressed in the EEC mitochondrial matrix. Therefore, by using this dual transgenic model, we can simultaneously measure EEC cytoplasmic Ca^2+^ levels and mitochondrial Ca^2+^ levels by measuring the change in green and red fluorescence (Video 4 and 5). We then analyzed how EEC cytoplasmic and mitochondrial Ca^2+^ levels change during development by tracing the same zebrafish from 3dpf to 6dpf (Fig. 5A-E’’). Our results showed that at 3dpf, the EECs exhibit low cytoplasmic and mitochondrial Ca^2+^ levels (Fig. 5B-B’’, G, H). However, at 4dpf, there is a significant increase in both EEC cytoplasmic and mitochondrial Ca^2+^ levels (Fig. 5C-C’’, G, H). From 5dpf to 6dpf, the EEC cytoplasmic Ca^2+^ levels decreased, while mitochondrial Ca^2+^ levels remained high (Fig. 5D-E’’, G, H). As a result, from 3dpf to 6dpf, the EEC mitochondrial to cytoplasmic Ca^2+^ ratio continues to increase (Fig. 5F). A similar trend was found in all of the zebrafish samples that we traced (Fig. 5I-K). When we grouped data of zebrafish of the same age together, we also observed that as the EECs became more mature, the EECs increased their mitochondrial to cytoplasmic Ca^2+^ level ratio (Fig. 5I-K).

**Figure 5.**
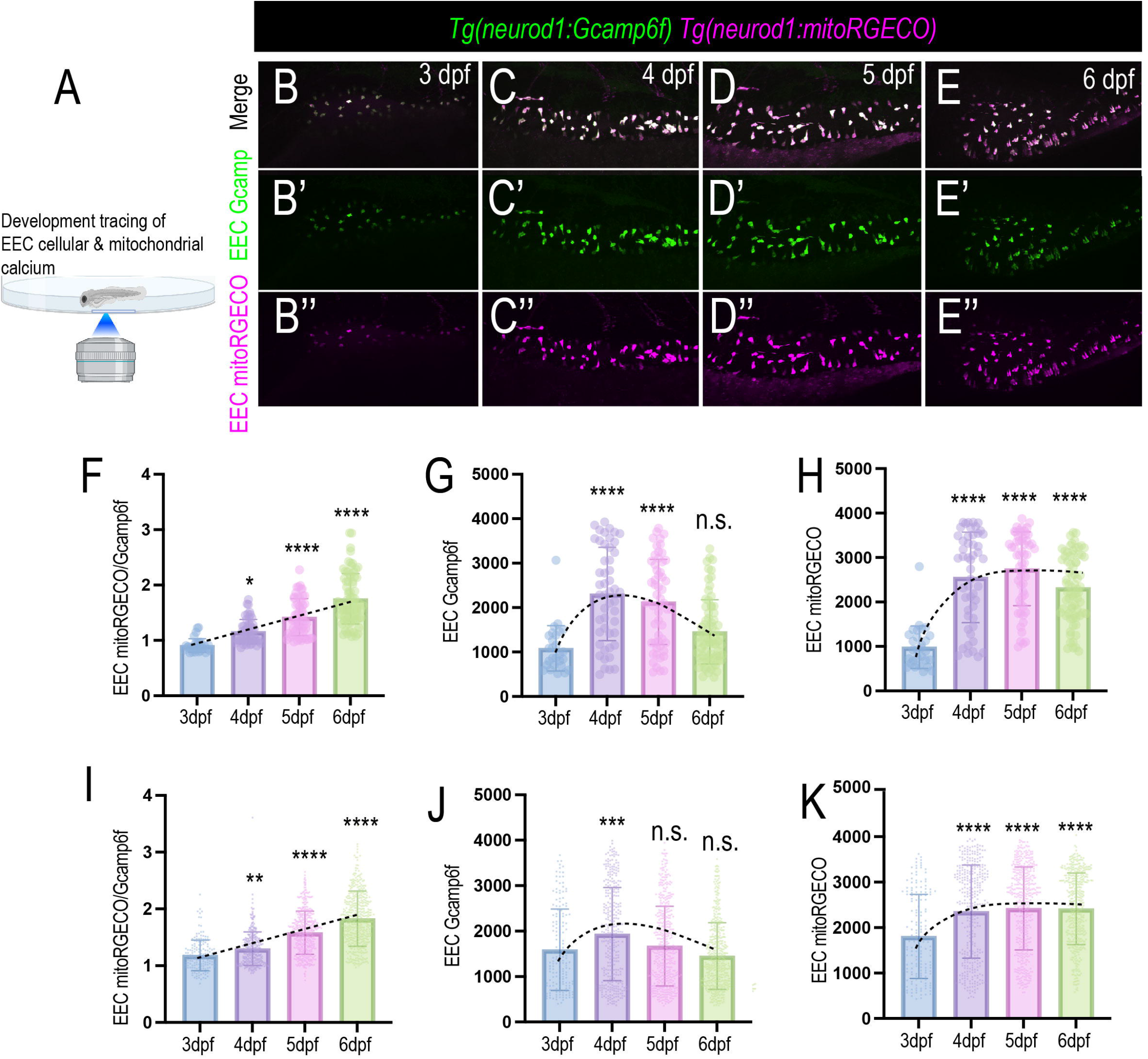
The change of EEC mitochondrial activity during development. (A) In vivo imaging to trace the EEC cytoplasmic and mitochondrial calcium in the same zebrafish from 3dpf to 6dpf. (B-E’’) Confocal projections of the same *Tg(neurod1:Gcamp6f); Tg(neurod1:mitoRGECO)* zebrafish at 3dpf, 4dpf, 5dpf, and 6dpf. The EEC cytoplasmic Ca^2+^ level is represented via the Gcamp6f fluorescence (green). The EEC mitochondrial Ca^2+^ level is displayed through mitoRGECO fluorescence (magenta). (F-H) Quantification of the EEC mitochondria-to-cytoplasmic Ca^2+^ ratio, cytoplasmic Ca^2+^, and mitochondrial Ca^2+^ in the zebrafish represented in B-E’’. (I-K) Combined quantification of 10 zebrafish EECs mitochondria-to-cytoplasmic Ca^2+^ ratio, cytoplasmic Ca^2+^, and mitochondrial Ca^2+^. Each dot in F-K represents an individual EEC. One-Way Anova followed by Tukey’s post test was used in F-K for statistical analysis. *p<0.05, ** p<0.01, *** p<0.001, **** p<0.0001.

### Gut microbiota increases resting EEC mitochondrial activity and spontaneous firing

Our new genetic zebrafish model and imaging approaches allow us to investigate how gut microbiota changes EEC cytoplasmic and mitochondrial activity in vivo. We generated *Tg(neurod1:Gcamp6f); Tg(neurod1:mitoRGECO)* GF and CV zebrafish and imaged the fish’s proximal intestinal EECs at 7dpf (Fig. 6A). First, we examined the absolute EEC cellular plasma Ca^2+^ levels and EEC mitochondrial Ca^2+^ levels by analyzing the individual EEC Gcamp6f and EEC mitoRGECO fluorescent levels in GF and CV zebrafish. Compared to GF EECs, CV EECs exhibit significantly lower cytoplasmic and mitochondrial Ca^2+^ levels (Fig. 6A-E). However, the CV EECs exhibit a significantly higher mitochondrial to cytoplasmic Ca^2+^ ratio (Fig. 6A-D, F). Moreover, many of the EECs in the CV but not GF zebrafish exhibit higher mitochondrial Ca2+ levels near the basal membrane (Fig. 6C-D). These results suggest that gut microbiota may promote low resting EEC cytoplasmic Ca^2+^ levels but enhance EEC mitochondrial activity. This finding is consistent with the results from our RNA sequencing analysis above (Fig.2).

**Figure 6.**
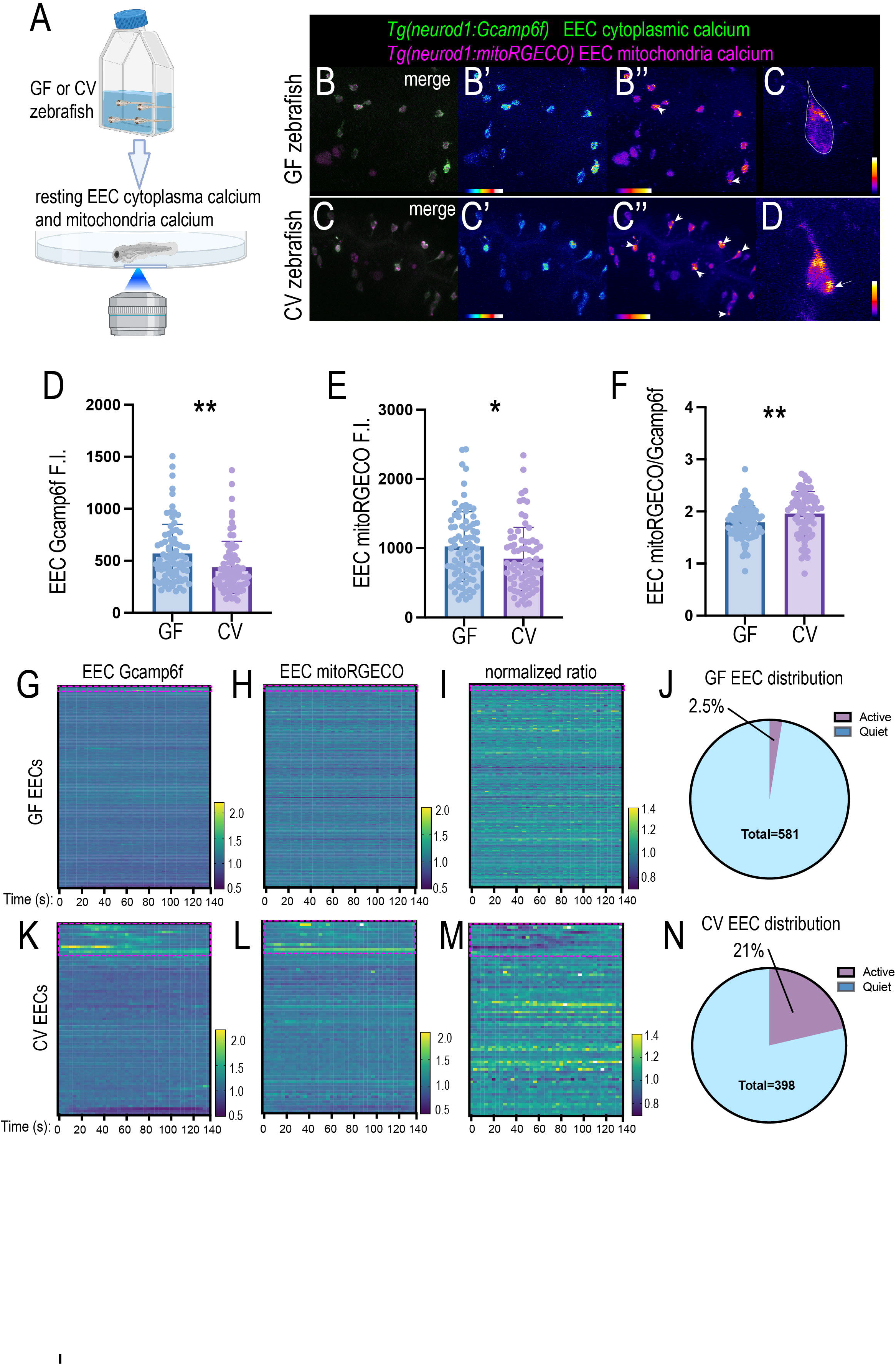
Commensal microbiota colonization alters the resting EEC cytoplasm and mitochondria calcium activity. (A) In vivo imaging to analyze the 7dpf GF and CV zebrafish EEC cellular and mitochondrial activity. (B-C’’) Confocal projection of the GF and CV *Tg(neurod1:Gcamp6f); Tg(neurod1:mitoRGECO)* zebrafish. Note that the absolute EEC Gcamp fluorescence is relatively higher in GF EECs. The EEC mitoRGECO/Gcamp ratio is lower in GF EECs. The white arrows in B’’ and C’’ indicates the EECs with higher mitochondrial activity near the base membrane. (C-D) zoom in view shows representative EECs in GF (C) and CV (D) zebrafish mitochondrial calcium activity. The CV EEC in D displayed high mitochondrial calcium near the base membrane (white arrow). (D-F) Quantification of absolute Gcamp, mitoRGECO, and mitoRGECO/Gcamp ratio in GF and CV zebrafish proximal intestinal EECs. Each dot represents an EEC. More than five zebrafish were analyzed in each condition. (G-M) Analyze the relative EEC Gcamp, EEC mitoRGECO, and EEC mitoRGECO/Gcamp ratio in GF and CV zebrafish on a temporal scale. The EEC Gcamp, EEC mitoRGECO, and EEC mitoRGECO/Gcamp ratio at each time point were normalized to t0. These quantification data sets reveal the dynamic change of EEC cellular and mitochondrial activity in GF and CV conditions. Each line in G-M represents an individual EEC. The red circles in G-M indicate the EECs that exhibit Gcamp fluorescence fluctuation. These EECs that display dynamic Gcamp fluorescence fluctuation were also referred to as active EECs in J and N. (J-N) Quantification of the percentage of the quiet and active EECs in GF and CV zebrafish. Student T-test was used in F for statistical analysis. * P<0.05, ** P<0.01.

Using a 3D cell tracking approach, we can automatically track individual EECs and analyze their Gcamp6f and mitoRGECO fluorescent change on a temporal scale. We analyzed the relative EEC cytoplasmic Ca^2+^ and mitochondrial Ca^2+^ levels change in GF and CV zebrafish. For each EEC, we normalized the EEC Gcamp6f, EEC mitoRGECO, and EEC mitoRGECO/Gcamp6f ratio value to their values at time 0. Our results indicate that some EECs in the CV zebrafish exhibit low amplitude firing as reflected by the temporal fluctuation of the EEC cytoplasmic Ca^2+^ levels (Fig. 6K-M) (Video 6). However, this spontaneous firing is not apparent in the GF zebrafish EECs (Fig. 6G-I) (Video 7). Analysis of EECs across different GF and CV zebrafish samples indicate that ∼21% of CV EECs exhibit low amplitude firing, but only 2.5% of GF EECs exhibit low amplitude firing (Fig. 6J, N). Those EECs with spontaneous firing increase the relative mitochondrial Ca^2+^ levels but not the relative mitochondria-to-cytoplasm Ca^2+^ ratio (Fig. 6H, L). These results suggest that at the resting condition, the CV EECs have higher mitochondrial activity, and their mitochondrial activity is more dynamic.

### Nutrient-induced EEC mitochondrial calcium increase requires gut microbiota

As the primary sensory cells, one of the major functions of EECs is to sense the intestinal lumen’s nutrients. Using the *Tg(neurod1:Gcamp6f)* zebrafish model, we previously developed a method to image live zebrafish’s EEC response to nutrients using an epifluorescent microscope [24]. Our previous methods allowed us to examine the systemic intestinal EEC nutrient response. However, it does not enable us to image the EECs’ nutrient sensing function at the cellular level [24]. To analyze how EECs respond to nutrients at the cellular level in live zebrafish, we developed a method to give stimulants to the zebrafish when we perform confocal imaging. Using 3D segmentation and 3D objective tracing, we can then quantify individual EECs’ response to nutrient stimulation systemically (Fig. 7A). Similar to mammals, the zebrafish EECs respond to nutrient stimulants like fatty acid [24, 38]. Previous studies suggest that long-chain fatty acids like linoleic acid bind with the G-protein coupled receptor in zebrafish EECs, stimulating calcium release from ER (Fig. 7B) [38]. Whether nutrient stimulation modulates EEC mitochondrial Ca^2+^ levels remains unclear.

**Figure 7.**
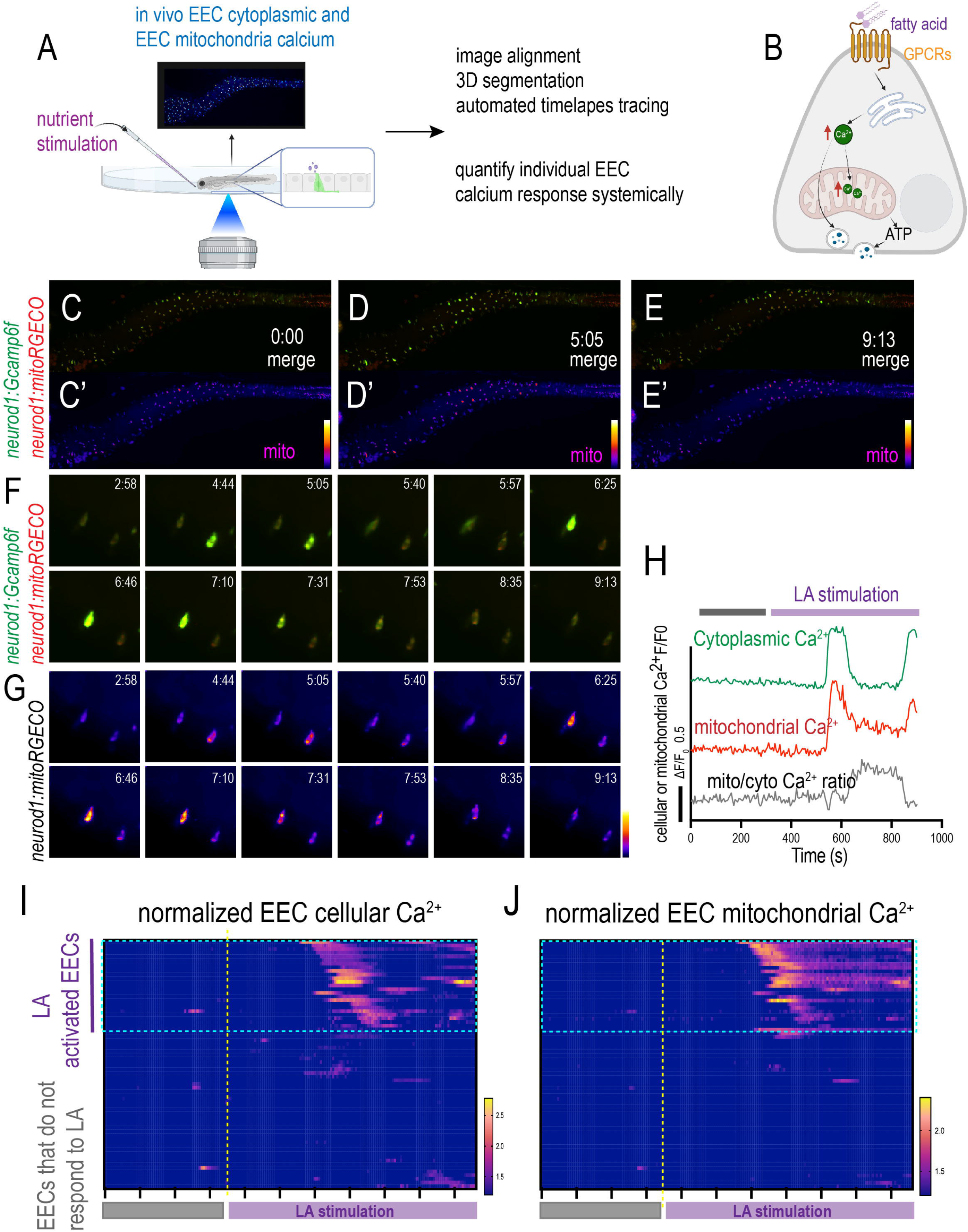
Analyze individual EEC cellular and mitochondrial activity in response to nutrient stimulation in live zebrafish. (A) Image and analyze the EEC cellular and mitochondrial calcium activity before and after nutrient stimulation using the confocal microscope. (B) The hypothesis model figure supported by our experimental data shows fatty acid increases both cytoplasmic and mitochondrial Ca^2+^. Long-chain fatty acids, such as linoleic acid (LA), binds the fatty acid receptor at the cell membrane. Activating the fatty acid receptor induces Ca^2+^ release from the ER and increases the cytoplasmic Ca^2+^. The increased cytoplasmic Ca^2+^ is then translocated into the mitochondrial matrix, which increases the mitochondrial matrix’s Ca^2+^ levels. The increased mitochondrial Ca^2+^ promotes ATP production and powers EEC vesicle secretion. (C-E’) Time-lapse images of the whole zebrafish intestinal EECs’ cytoplasmic and mitochondrial calcium change post linoleic acid stimulation. The EEC cytoplasmic Ca^2+^ was labeled by Gcamp6f (green), and the EEC mitochondrial Ca^2+^ was labeled by mitoRGECO (red). (F-G) Zoom-out view shows two representative EECs that are activated by linoleic acid. (H) Analysis of fluorescence change of Gcamp, mitoRGECO, and mitoRGECO/Gcamp ratio in a presentative linoleic acid activated EEC. (I-J) Analysis of fluorescence change of Gcamp and mitoRGECO of 68 EECs in one zebrafish before and after linoleic acid stimulation. The EECs that increase cytoplasmic Ca^2+^ were defined as “LA activated EECs”. Noted that the majority of the linoleic activated EECs also exhibit increased mitochondrial calcium.

Using in vivo EEC calcium imaging and 3D automated cell tracing, we measured the individual EECs cytoplasmic and mitochondrial calcium response to nutrient stimulation systemically in live zebrafish (Fig. 7A) (video 8,9). Our results demonstrated that nutrients such as linoleic acid stimulate a subset of EECs and increase those EECs’ cytoplasmic Ca^2+^ level (Fig. 7C-G) (Video 8, 9). Along with the EECs’ cytoplasmic calcium increase, there is also a consistent mitochondrial calcium increase following nutrient stimulation (Fig. 7C-G) (Video 8, 9). In the linoleic acid-activated EECs, linoleic stimulation induces cytoplasmic calcium peak in these cells and the cellular cytoplasmic Ca^2+^ levels returned to their basal activity level (Fig. 7F-H). Their mitochondrial Ca^2+^ levels increased immediately following the cytoplasmic Ca^2+^ peak (Fig. 7F-H). However, unlike the cytoplasmic Ca^2+^, the mitochondrial Ca^2+^ level remained higher than the basal Ca^2+^ level after the peak (Fig. 7F-H). As a result, the relative mitochondrial-to-cytoplasmic Ca^2+^ ratio increased post-linoleic acid stimulation (Fig. 7F-H). Moreover, our results demonstrate that the nutrient induced mitochondrial Ca^2+^ increase is more prominent in the mitochondria near the basal membrane (Fig. S4). This suggests that the nutrient induced mitochondrial Ca^2+^ increase is likely linked with the EECs’ vesicle secretion process. Our results further indicate that the majority of the linoleic acid activated EECs in the conventionally raised zebrafish exhibit elevated mitochondrial calcium in response to nutrient stimulation (Fig. 7I-J).

Finally, we investigated whether and how gut microbiota regulates the EECs’ nutrient response. We generated GF and CV *Tg(neurod1:Gcamp6f); Tg(neurod1:mitoRGECO)* zebrafish (Fig. 8A). We then stimulated the GF and CV zebrafish with linoleic acid and recorded how the GF and CV EECs responded to the linoleic stimulation. Our results indicate that, compared with CV zebrafish, the percentage of EECs that can be activated by linoleic acid in GF zebrafish is less (Fig. 8B). Within the activated EECs, the cytoplasmic Ca^2+^ amplitude remains the same between GF and CV groups (Fig. 8C). However, within the activated EECs, the mitochondrial Ca^2+^ amplitude significantly increased in the CV EECs (Fig. 8D, E-F). The same result is shown when we trace the temporal EEC cytoplasmic and mitochondrial Ca^2+^ levels in GF and CV zebrafish (Fig. 8G-L). In most CV EECs, nutrient stimulation activates both cytoplasmic and mitochondrial Ca^2+^ and increases the mitochondrial to cytoplasmic Ca^2+^ ratio (Fig. 8G). However, the nutrient induced EEC mitochondrial activation is significantly reduced in GF EECs (Fig. 8G-L). The nutrient induced mitochondrial-to-cytoplasmic Ca^2+^ ratio increase is also impaired in GF EECs (Fig. 8I). These results suggest that the nutrient-induced EEC mitochondrial activation requires signals from commensal microbiota colonization.

**Figure 8.**
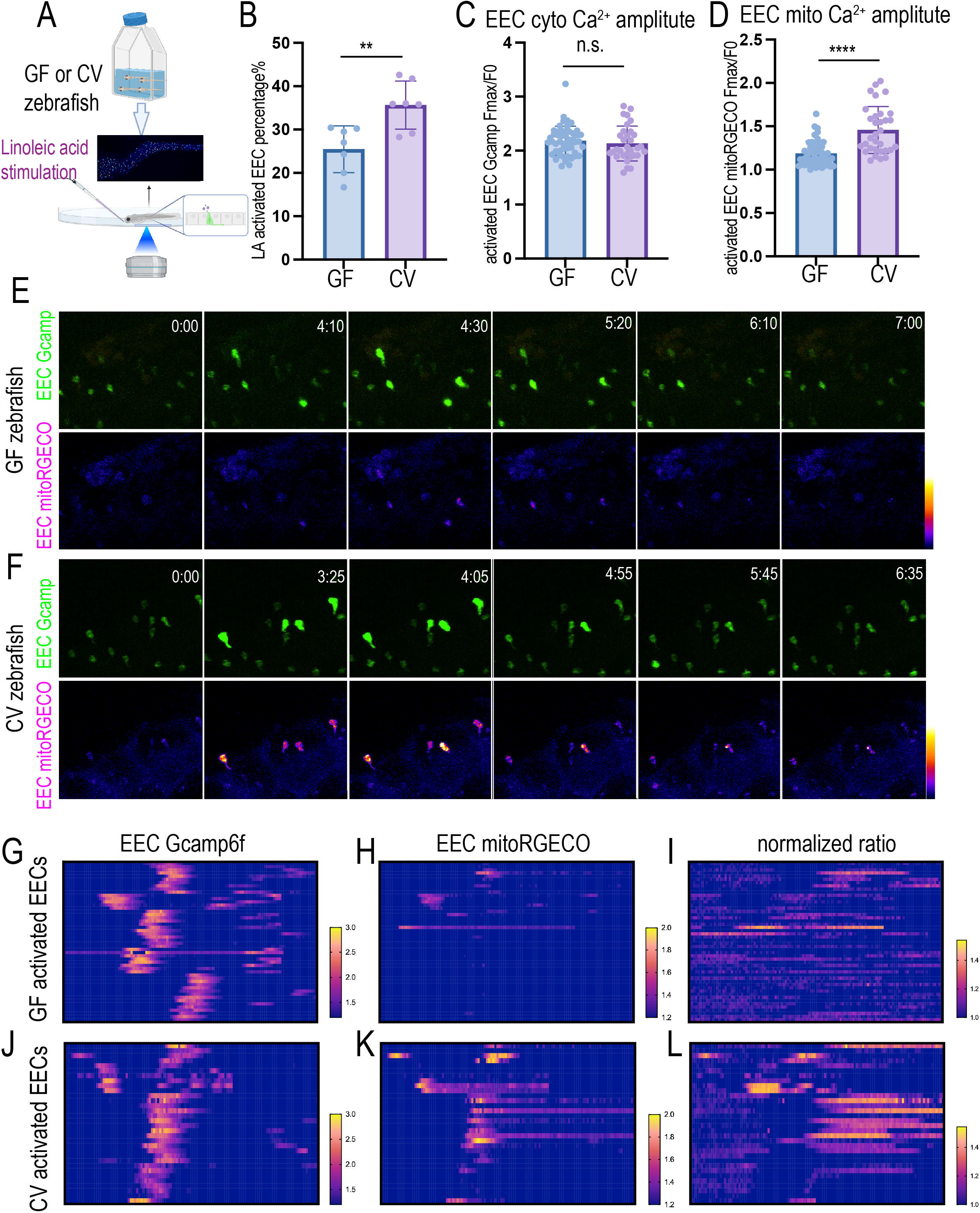
Nutrient induced EECs’ mitochondrial calcium increase requires commensal microbiota colonization. (A) In vivo imaging to analyze the 7dpf GF and CV zebrafish EEC cellular and mitochondrial activity in response to linoleic acid stimulation. (B) Quantification of the percentage of EECs that are activated by linoleic acid in GF and CV zebrafish. Each dot represents an individual zebrafish. (C-D) Quantification of the linoleic acid-activated EEC cytoplasmic Ca^2+^ amplitudes and the mitochondrial Ca^2+^ amplitudes in GF and CV zebrafish. Each dot represents an individual EEC. Note that there is no significant change in the cytoplasmic Ca^2+^ increase. However, CV EECs exhibit a significantly higher mitochondrial Ca^2+^ increase in response to linoleic acid stimulation. (E-F) Time-lapse images of the representative EECs post linoleic acid stimulation in GF and CV zebrafish. The EEC cytoplasmic Ca^2+^ was labeled by Gcamp6f, and the EEC mitochondrial Ca^2+^ was labeled by mitoRGECO. (G-L) Analysis of the change of the EEC Gcamp6f fluorescence, EEC mitoRGECO fluorescence, and EEC mitoRGECO/Gcamp6f ratio in GF and CV zebrafish. Only the linoleic acid activated EECs were plotted. G & J, H & K, and I & J used the same heatmap scale. 32 activated EECs from 4 CV zebrafish and 50 activated EECs from 5 GF zebrafish were analyzed.

## Discussion

In this study, by using transcriptomics, genetics, in vivo imaging, and gnotobiotic manipulation, we revealed that commensal microbiota colonization is critical in shaping EEC maturation and function during development (Fig. S5). Importantly, our data revealed that commensal microbiota colonization is essential in promoting mitochondrial activity and nutrient induced mitochondrial activation. Selectively manipulating gut microbial signals to alter EEC mitochondrial function may open new opportunities to change EEC vesicle secretion and EEC-neuronal communication.

### The change of EEC mitochondrial activity during development

Using in vivo imaging to track EECs during development in live zebrafish, our results revealed that EEC mitochondrial activity is dynamically regulated during development. Our data showed shortly after commensal microbiota colonization, EECs increase both cytoplasmic and mitochondrial calcium activity. A phenomenon we referred to as the “EEC awakening”. After the EEC awakening, the EECs down-regulate their cytoplasmic calcium levels but up-regulate their mitochondrial-to-cytoplasmic Ca^2+^ ratio. As sensory cells, it is critical for EECs to maintain low cytoplasmic Ca^2+^ levels to enable a depolarization potential. When EECs sense nutrient stimulants, the calcium ion channel on the cell membrane or ER membrane will open [2]. Ca^2+^ from the extracellular space or the ER lumen will flux into the cytoplasm matrix following the Ca^2+^ gradient. The low cytoplasmic Ca^2+^ levels are, therefore, essential to generating the gradient to produce the Ca^2+^ peak to trigger the downstream cellular signaling events [39]. Maintaining the membrane potential or the low cytoplasmic Ca^2+^ levels consumes ATP [39]. ATP can be generated via glycolysis or through the tricarboxylic acid cycle (TCA cycle) and oxidative phosphorylation mediated by mitochondria [40]. It is well appreciated that mitochondrial and metabolic remodeling is a central feature of differentiation and reprograming events [17]. The mitochondrial oxidative metabolism is often suppressed in stem cells [41, 42]. The stem cells, including intestinal stem cells, rely on glycolysis to generate ATP [43, 44]. The mitochondria in stem cells remain functional [43]. However, stem cells possess multiple mechanisms to suppress mitochondrial activity [44–46]. Upon differentiation, mitochondrial activity increases [43, 44]. On the one hand, the increased mitochondrial activity fuels the high metabolic demand of the differentiated cells. On the other hand, the increased mitochondrial activity generates necessary signaling molecules such as reactive oxygen species (ROS) and biosynthetic metabolites through the TCA cycle to promote the differentiation process [47, 48]. The mitochondria-derived signaling molecules also promote epigenetic remodeling and modulate gene expression [49]. Using an *in vivo* EEC mitochondria imaging approach to trace the same zebrafish across different developmental time points, our results revealed that the immature EECs at 3 days post fertilization display low mitochondrial activity. When zebrafish develop and EECs start to be functional, mitochondrial activity increases. The increased mitochondrial activity may not only provide energy to fuel the EEC cellular process but also provide the signaling that is necessary for the EECs to mature and function. Interestingly, in addition to the change in mitochondrial activity, we also observed changes in intracellular mitochondrial distribution during development. Specifically, our results revealed the increased mitochondrial contents at the basal lateral membrane during the EEC maturation process. The increased mitochondrial distribution in the mature EECs is likely to match the ATP demand of the vesicle secretion process in the basal lateral membrane.

In addition to providing energy, mitochondria also function as an important calcium buffer. In response to extracellular stimulation, cytoplasmic Ca^2+^ levels increase. This increase in cytoplasmic Ca^2+^ is quickly dissipated into intracellular organelles, such as the ER or mitochondria. In most cells, mitochondrial calcium uptake is mediated by the mitochondrial calcium uniporter (MCU), a calcium transporter protein in the mitochondrial inner membrane. The electrochemical potential across the mitochondrial inner membrane, generated by the respiration chain reaction, is the major driving force that enables calcium influx into the mitochondrial matrix via MCU. Cytoplasmic and mitochondrial Ca^2+^ coupling have not been studied in EECs. Our studies revealed that, in response to nutrient stimulation, a subset of EECs increase cytoplasmic Ca^2+^ activity. Following the increase of cytoplasmic Ca2+ levels, the mitochondrial Ca^2+^ levels increase in most activated EECs. Though the cytoplasmic Ca^2+^ quickly returned to the basal level, most EEC mitochondrial Ca^2+^ was continuously maintained at a high level. Basal mitochondrial respiratory function might be the key to mediating Ca^2+^ flux into the mitochondrial matrix. The increase in mitochondrial Ca^2+^ will increase mitochondrial respiration to sustain the ATP production that is required for the EEC vesicle secretion in response to the nutrient stimulants [20]. Our results show that mitochondria are concentrated near the basal membrane, where vesicle secretion occurs. Nutrient-induced mitochondrial activation is also most prominent in the mitochondria near the base membrane. This evidence supports the hypothesis that mitochondrial activation assists with vesicle secretion in mature EECs.

### The change of EEC morphology during development

In addition to the change in EEC mitochondrial activity, another hallmark of EEC maturation, revealed by our study, is the change in EEC morphology. Our study illustrated for the first time that immature EECs possess dynamic and active actin filaments in the basal membrane. However, the actin filaments disappeared in mature EECs. Instead, some mature EECs formed an elongated basal lateral membrane process, a structure that resembles the “neuropod” reported in previous mammalian studies [3]. Previous studies demonstrated that the neuropod structure enriches the neurofilaments and mitochondria [50]. EECs use neuropods to form synaptic connections with the underlying nerve terminates, including the vagal sensory nerve [3, 4]. What regulates the EEC neuropod formation and guides the EEC-neuronal synaptic connection remains unknown. In developing neurons, neurites form actin-supported extensions known as growth cones which seek synaptic targets [51]. Formation of the pre- and post-synaptic structures disables the filipodium-enriched actin structure at the leading age [52]. Can the EECs form a growth cone-like structure to find their targets and form synaptic connections with the neurons? Our results revealed that the immature EECs form thin actin-based elements in the basal lateral membrane, a structure that is similar to the filopodia projections found in the developing neuron axon growth cone. In the zebrafish that are colonized with commensal microbiota, these thin actin filaments in some of the EECs are replaced by the “neuropod-like” structure when EECs mature. This morphology evidence supported the hypothesis that the immature EECs may use active actin filaments to find the synaptic targets and form the synaptic connections with the underlying neurons. Establishing the EEC-neuronal connection will facilitate the EECs to transmit the ingested nutrients to the nervous system.

### How does gut microbiota regulate EEC maturation and mitochondrial function?

A major finding revealed by our study is that commensal microbiota colonization is critical in supporting EEC maturation and promoting EEC mitochondrial function. Our results established that microbiota colonization during early development might be critical in establishing the organisms’ appropriate nutrient-sensing function via promoting EEC maturation. Our results also suggested that postnatal microbiota colonization might be critical in promoting the formation of EEC-vagal neuronal communication as the commensal microbiota colonization promotes the formation of the neuropod-like structure. The EEC-vagal neuronal connection is essential in mediating gut-brain nutrient sensing. Gut microbiota can therefore modulate EEC or EEC-vagal communication to regulate brain nutrient perception and feeding behavior. Our results revealed that the EECs remain in an immature state and exhibit low mitochondrial activity when commensal microbiota is absent. Disrupting the commensal microbiota colonization or inhibiting the formation of the healthy postnatal microbiome may produce devasting effects on gut nutrient perception and metabolic regulation. The formation of the postnatal gut microbial community is influenced by many factors (maternal microbiome, delivery method, milk-feeding vs formula feeding). Previous research showed that disrupting the infant microbiome through antibiotic exposure results in many side effects, including obesity and weight gain later in life [53]. The EECs are critical in sensing ingested nutrients and maintaining homeostasis [2]. Our study suggests that disrupting the commensal microbial community early in life will change the EECs’ function and maturity, which may change how the body responds to ingested nutrients and affect energy homeostatic control.

The mitochondrial energetic adaptations encompass a conserved process that maintains cell and organisms’ fitness in the changing environment [54]. Our studies suggested that in response to the commensal microbiota colonization, the EECs’ increase mitochondrial respiration and enhance the mitochondrial calcium activity. Our transcriptomic data revealed that microbial induced EECs’ energy and mitochondrial adaptation are involved with the increased mitochondrial cristae formation and increased mitochondrial respiratory chain assembly via enhancing mitochondrial protein import and facilitating protein translation in the mitochondrial matrix (Fig. 2I). The EECs’ mitochondrial energetic adaptation in response to commensal microbiota colonization may contribute to the systemic host adaptation to microbial colonization that is to compete to the limited nutrients, enhance nutrient utilization efficiency and promote nutrient storage. The microbial and molecular mechanisms by which microbial signals regulate mitochondrial activity and intercede with the nutritional metabolism pathway within the EECs are intriguing questions that require future investigations.

## Acknowledgments

This work was supported by NIH K01-DK125527, The OSU Food for Health Research Initiative Innovation Seed Grant, and The Global Probiotic Young Investigator Grant.

## Author contributions

Alfahadh Alsudayri conducted the gnotobiotic experiments, some immunofluorescence staining, and performed most of the data analysis. Shane Perelman performed the experiments for EEC mitochondrial temporal tracking and data analysis. Annika Chura performed some immunofluorescence staining experiments. Melissa Brewer and Maddie McDevitt contributed to the data analysis and facilitated the experiments. Amrita Mandal generated the *Tg(neurod1:mitoRGECO)* and the *Tg(neurod1:mitoEOS)* transgenic zebrafish models. Dr. Catherine Drerup provided the *Tg(neurod1:mitoRGECO)* and the *Tg(neurod1:mitoEOS)* transgenic zebrafish models and provided technique and conceptual instructions. Lihua Ye directed the project and wrote the manuscript.

## Methods

### Zebrafish strains and husbandry

All zebrafish experiments conformed to the US Public Health Service Policy on Humane Care and Use of Laboratory Animals, using protocol number 2021A00000091 approved by the Institutional Animal Care and Use Committee of the Ohio State University. Conventionally-reared adult zebrafish were reared and maintained on a recirculating aquaculture system using established methods [28]. For experiments involving conventionally-raised zebrafish larvae, adults were bred naturally in system water and fertilized eggs were transferred to 100mm petri dishes containing ∼25mL of egg water at approximately 6 hours post-fertilization. The resulting larvae were raised under a 14h light/10h dark cycle in an air incubator at 28°C at a density of 2 larvae/ml water. All the experiments performed in this study ended at 7dpf unless specifically indicated. The zebrafish lines used in this study are listed in Table 2. All lines were maintained on a EKW background.

### Gnotobiotic zebrafish husbandry

For experiments involving gnotobiotic zebrafish, we used our established methods to generate germ-free zebrafish using natural breeding (ref.) with the following exception: Gnotobiotic Zebrafish Medium (GZM) with antibiotics (AB-GZM) was supplemented with 50 μg/ml gentamycin (Sigma, G1264). Germ-free zebrafish eggs were maintained in cell culture flasks with GZM at a density of 1 larvae/ml. From 3 dpf to 7 dpf, 60% daily media change, and newborn fish food (Ultra Fresh Ltd.) feeding were performed as described [28].

To generate conventionalized zebrafish, 15 mL filtered system water (5μm filter, SLSV025LS, Millipore, final concentration of system water ∼30%) was inoculated to flasks containing germ-free zebrafish in GZM at 3 dpf when the zebrafish normally hatch from their protective chorions. The same feeding and media change protocol was followed as for germ free zebrafish. Microbial colonization density was determined via Colony Forming Unit (CFU) analysis. To analyze the effect of high fat feeding on intestinal bacteria colonization, dissected digestive tracts were dissected and pooled (5 guts/pool) into 1mL sterile phosphate buffered saline (PBS) which was then mechanically disassociated using a Tissue-Tearor (BioSpec Products, 985370). 100 µL of serially diluted solution was then spotted on a Tryptic soy agar (TSA) plate and cultured overnight at 30°C under aerobic conditions.

### Zebrafish EEC RNA sequencing analysis

The zebrafish EEC RNA sequencing data was generated in our previous study (ref.). This dataset can be accessed at GSE151711. Conventionalized (CV) and germ-free (GF) *TgBAC(cldn15la:EGFP); Tg(neurod1:TagRFP)* ZM000 fed zebrafish larvae were derived and reared using the published protocol [28] for Flow Activated Cell Sorting (FACS) to isolate zebrafish EECs and other IECs. The protocol for FACS was adopted from a previous publication [27]. Replicate pools of 50-100 double transgenic *TgBAC(cldn15la:EGFP); Tg(neurod1:TagRPF)* zebrafish larvae were euthanized with Tricaine and washed with deyolking buffer (55 mM NaCl, 1.8 mM KCl and 1.25 mM NaHCO3) before they were transferred to dissociation buffer [HBSS supplemented with 5% heat-inactivated fetal bovine serum (HI-FBS, Sigma, F2442) and 10 mM HEPES (Gibco, 15630–080)]. Larvae were dissociated using a combination of enzymatic disruption using Liberase (Roche, 05 401 119 001, 5 μg/mL final), DNaseI (Sigma, D4513, 2 μg/mL final), Hyaluronidase (Sigma, H3506, 6 U/mL final) and Collagenase XI (Sigma, C7657, 12.5 U/mL final) and mechanical disruption using a gentleMACS dissociator (Miltenyi Biotec, 130-093-235). 400 μL of ice-cold 120 mM EDTA (in 1x PBS) wwas added to each sample at the end of the dissociation process to stop the enzymatic digestion. Following addition of 10 mL Buffer 2 [HBSS supplemented with 5% HI-FBS, 10 mM HEPES and 2 mM EDTA], samples were filtered through 30 μm cell strainers (Miltenyi Biotec, 130-098-458). Samples were then centrifuged at 1800 rcf for 15 minutes at room temperature. The supernatant was decanted, and cell pellets were resuspended in 500 μL Buffer 2. FACS was performed with a MoFlo XDP cell sorter (Beckman Coulter) at the Duke Cancer Institute Flow Cytometry Shared Resource. Single-color control samples were used for compensation and gating. Viable EECs or IECs were identified as 7-AAD negative.

Samples from three independent experimental replicates were performed. 250-580 EECs (n=3 for each CV and GF group) and 100 IECs (n=3 for each CV and GF group) from each experiment were used for library generation and RNA sequencing. Total RNA was extracted from cell pellets using the Argencourt RNAdvance Cell V2 kit (Beckman) following the manufacturer’s instructions. RNA amplification prior to library preparation had to be performed. The Clontech SMART-Seq v4 Ultra Low Input RNA Kit (Takara) was used to generate full-length cDNA. mRNA transcripts were converted into cDNA through Clontech’s oligo(dT)-priming method. Full length cDNA was then converted into an Illumina sequencing library using the Kapa Hyper Prep kit (Roche). In brief, cDNA was sheared using a Covaris instrument to produce fragments of about 300 bp in length. Illumina sequencing adapters were then ligated to both ends of the 300bp fragments prior to final library amplification. Each library was uniquely indexed allowing for multiple samples to be pooled and sequenced on two lanes of an Illumina HiSeq 4000 flow cell. Each HiSeq 4000 lane could generate >330M 50bp single end reads per lane. This pooling strategy generated enough sequencing depth (∼55M reads per sample) for estimating differential expression. Sample preparation and sequencing was performed at the GCB Sequencing and Genomic Technologies Shared Resource.

Zebrafish RNA-seq reads were mapped to the danRer10 genome using HISAT2(Galaxy Version 2.0.5.1) using default settings. Normalized counts and pairwise differentiation analysis were carried out via DESeq2. The significance threshold of p < 0.05 was used for comparison.

### Immunofluorescence staining

Whole mount immunofluorescence staining was performed as previously described [24]. In brief, ice cold 2.5% formalin was used to fix zebrafish larvae overnight at 4°C. The samples were then washed with PT solution (PBS+0.75%Triton-100). The skin and remaining yolk were then removed using forceps under a dissecting microscope. The deyolked samples were then permeabilized with methanol for more than 2 hrs at −20°C. Samples were then blocked with 4% BSA at room temperature for more than 1 hr. The primary antibody was diluted in PT solution and incubated at 4°C for more than 24 hrs. Following primary antibody incubation, the samples were washed with PT solution and incubated overnight with secondary antibody with Hoechst 33342 for DNA staining. Imaging was performed with Nikon AXR confocal using the 20× of 40× water immersion lens. The primary antibodies were listed in Table 1. The secondary antibodies in this study were from Alexa Fluor Invitrogen were used at a dilution of 1:250.

### Live imaging and image analysis

The zebrafish larvae were anesthetized with Tricaine methanesulfonate (MS222) and were mounted in the 35mm confocal dish using 1% low-melting-Agar. All the in vivo imaging were performed using the Nikon AXR confocal. When imaging the EEC cellular and mitochondrial calcium activity using the *Tg(neurod1:Gcamp6f); Tg(neurod1:mitoRGECO)* zebrafish, the zebrafish were not anesthetized due to the effects of Tricaine in activating EECs. In the developmental tracing experiments, after imaging, the zebrafish were dug out of the Agar, placed in 6-well plate, and returned to the incubator until the next imaging time point. In the experiments when the temporal EEC activity was traced, the images were collected using the resonate scanner. It takes less than 10 seconds to collect the whole intestinal z-stack. The interval of time frames is 10 seconds. In the experiments when the nutrient stimulants were applied. A small window was cut in front of the zebrafish, which allowed the zebrafish mouth to be exposed. First, the zebrafish intestine was imaged before the stimulants were applied to assess the basal line EEC activity. After collecting the baseline EEC activity, the image acquisition was pulsed, and nutrient stimulants were added. The egg water in the confocal dish was removed. 1ml nutrient stimulate solution was delivered into the window in front of the zebrafish. After the nutrient stimulation was applied, the image acquisition process resumes. The time lapse images were collected to assess the nutrient induced EEC activation. For the image analysis to assess the EEC calcium activity, the images were first aligned using the Nikon NLS element software. The threshold was defined using the Gcamp channel to perform segmentation of the individual EECs and identify individual EEC units. The non-EECs were filtered out via the shape criteria, the fluorescence intensity, and the size. The individual EEC units in different time frames were traced and tracked via the NLS element 3D-object tracing software. Due to the issues of gut motility, not every EEC in the zebrafish can be successfully traced throughout the time course. The mean fluorescence intensity of the individual EEC in each time frame will be calculated. The cluster 3.0 software was used to perform the clustering analysis of the EECs that exhibit different temporal calcium dynamics.

## Supplemental Figure Legends

**Supplemental Figure 1.**
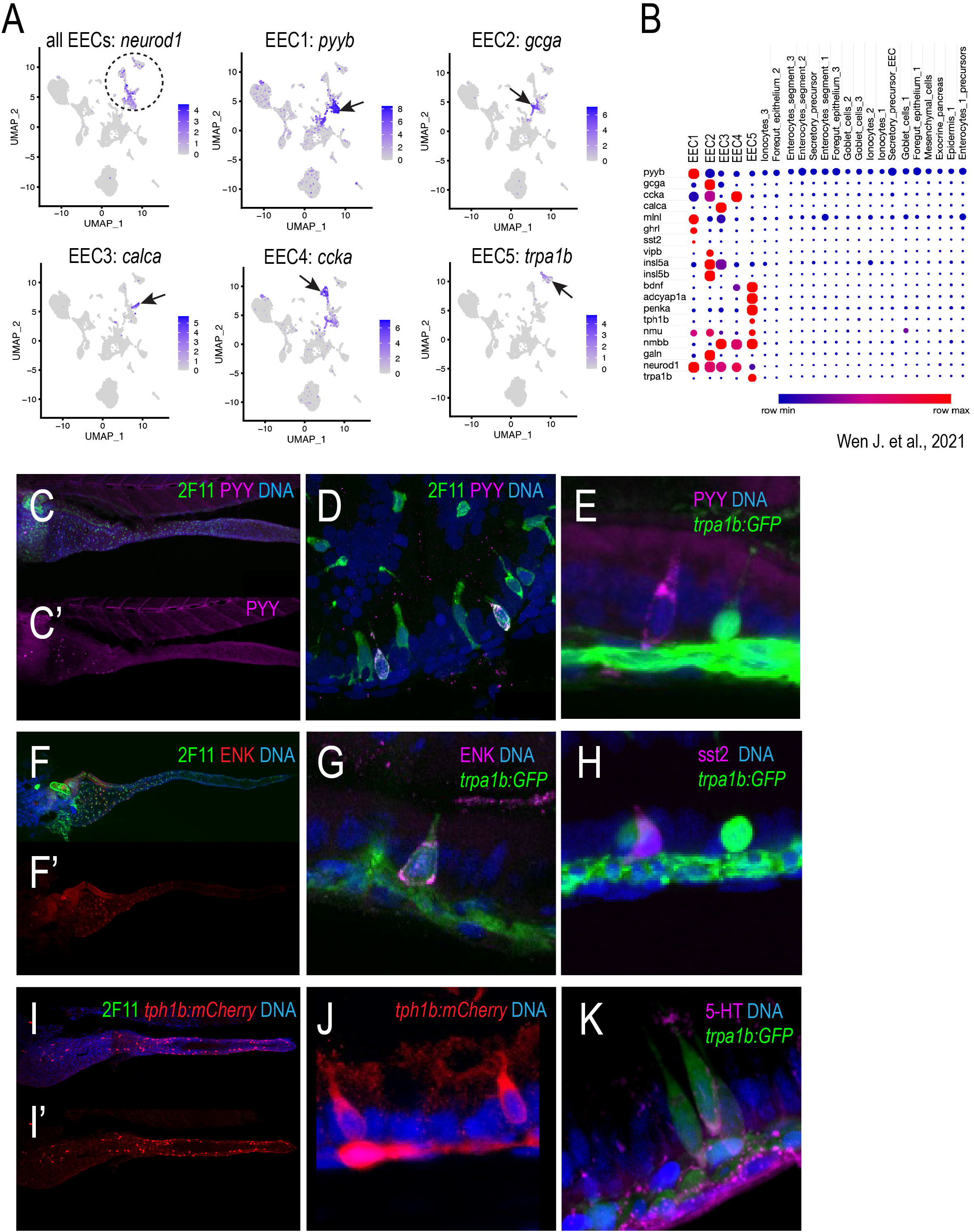
The EEC subtypes in zebrafish larvae. (A) UMAP plots of the zebrafish intestine single-cell RNA sequencing showing the zebrafish EECs and the five EEC subtypes in zebrafish larvae. The zebrafish scRNA dataset is from Wen J. et al., 2021. (B) The hormone profiles in the five zebrafish EEC subtypes. (C-E) Immunofluorescence staining of the PYY+EEC subtype. Note that the PYY+EECs are distributed in the proximal zebrafish intestine (C-C’). It overlaps with the secretory cell marker 2F11 (D) but does not overlap with the marker for other EEC subtypes, such as trpa1b (E). (F-K) Immunofluorescence staining of the Trpa1+EEC subtype. The single-cell RNA seq data above demonstrate that the Trpa1+EECs (EEC5) express the peptide enkephalin (ENK) and the enzyme that synthesizes serotonin (tph1b). (F-G) Immunofluorescence staining of ENK confirms that only Trpa1+EECs express ENK (G). Interestingly, ENK is only expressed in the Trpa1+EECs in the proximal intestine (F-F’). (H) Trpa1+EECs do not express sst2, a marker for the EEC subtype 1. (I-J) Tph1b is expressed in the EECs. (K) Immunofluorescence staining showing part of the Trpa1+EECs express 5-HT.

**Supplemental Figure 2.**
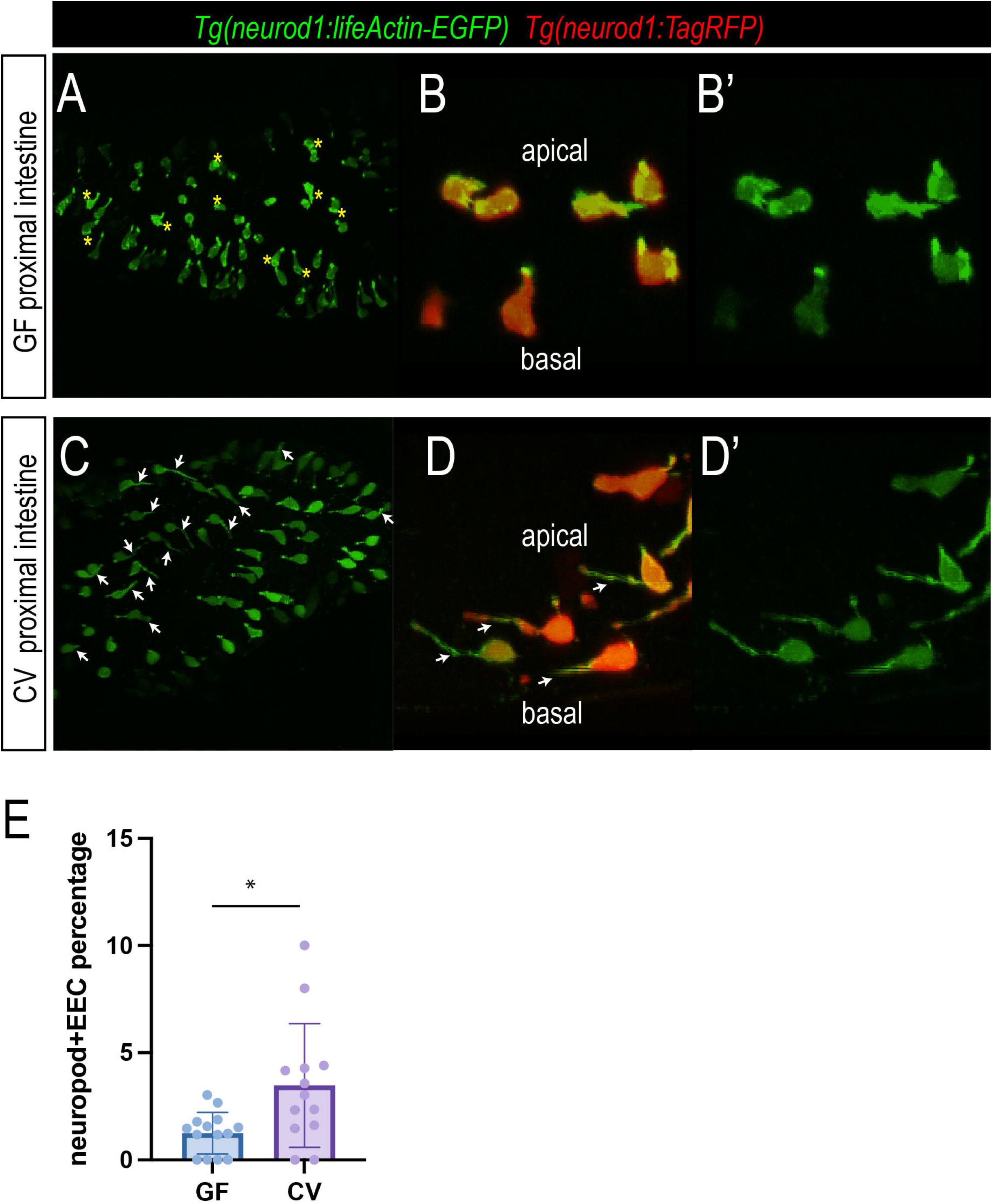
Commensal microbiota colonization promotes the formation of “neuropod”-like structure in EECs. (A-D’) Confocal projections of the GF and CV *Tg(neurod1:lifeActin-EGFP)* zebrafish. The yellow stars in A indicate EECs with thin actin filaments at the basal lateral membrane. The White arrows in C and D indicate the “neuropod” like elongated basal lateral membrane protrusions in CV EECs. (E) Quantification of the EEC percentage that has “neuropod” like structure in GF and CV conditions. Student T-test was used in E. Each dot represents an individual zebrafish. * P<0.05.

**Supplemental Figure 3.**
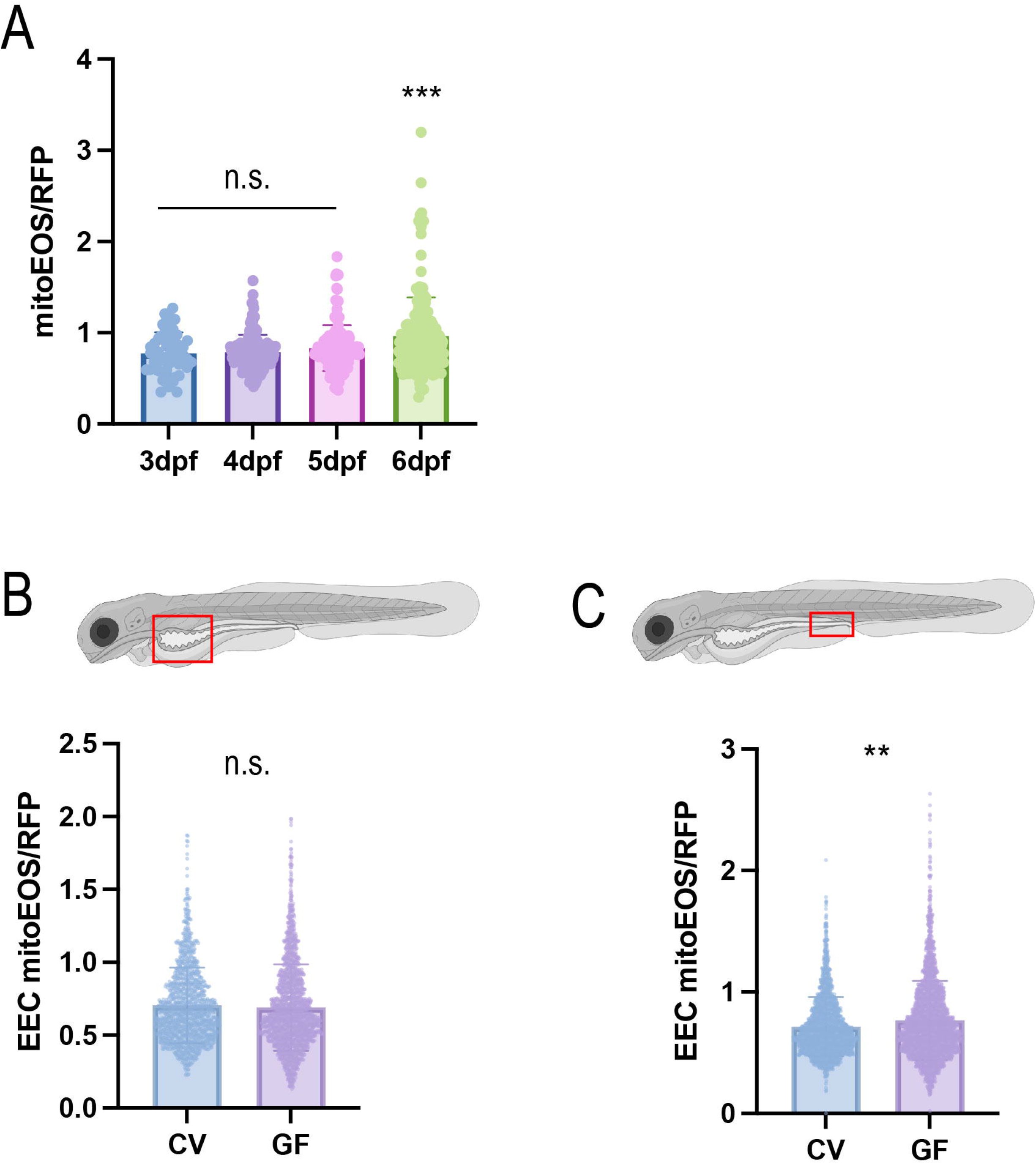
Gut microbiota did not alter the proximal intestine mitochondria abundance. (A) Quantification of the intracellular mitochondria abundance in the 3dpf to 6dpf zebrafish EECs. The mitochondrial abundance is represented by the *neurod1:mitoEOS* and *neurod1:RFP* fluorescence ratio in individual EECs. Each dot represents an EEC. 4 zebrafish were analyzed. (B-C) Quantification of the intracellular mitochondria abundance in the proximal and distal intestines of the GF and CV zebrafish. The mitochondrial abundance is represented by the *neurod1:mitoEOS* and *neurod1:RFP* fluorescence ratio in individual EECs. Each dot represents an EEC. More than 7 zebrafish in GF and CV groups were quantified. One-Way Anova followed by Tukey post-test was used in A. Student T-test was used in B and C. *** P<0.001, ** P<0.01.

**Supplemental Figure 4.**
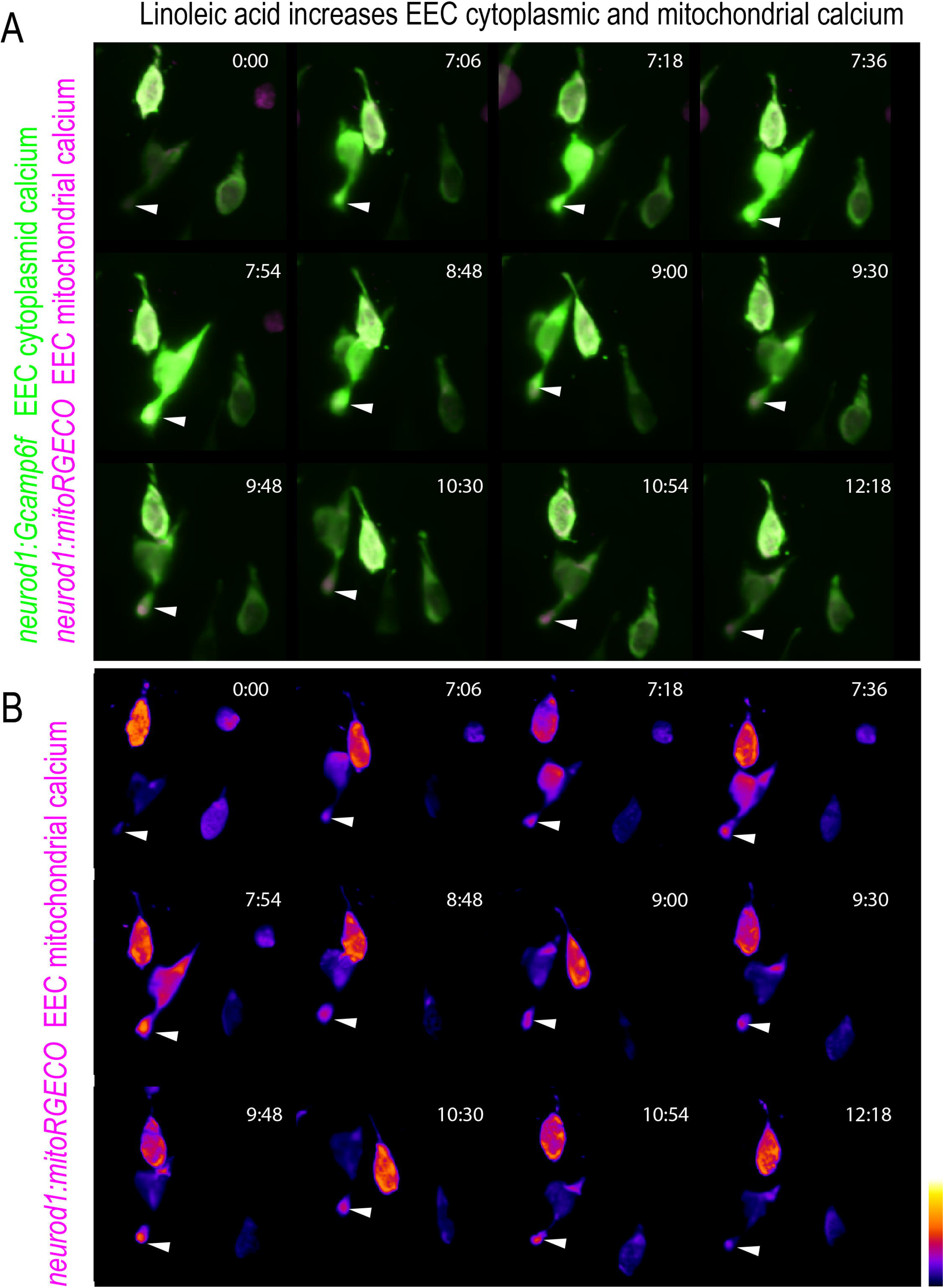
Nutrient stimulation prominently promotes mitochondrial Ca2+ to arise near the basal membrane. (A-B) Zoom in view shows two linoleic acid activated EECs in *Tg(neurod1:Gcamp6f); Tg(neurod1:mitoRGECO)* zebrafish. The cytoplasmic calcium was indicated with Gcamp6f (green fluorescence), and the mitochondrial calcium was indicated with mitoRGECO (magenta fluorescence). White arrows show activated mitochondria near the basal membrane.

**Supplemental Figure 5.**
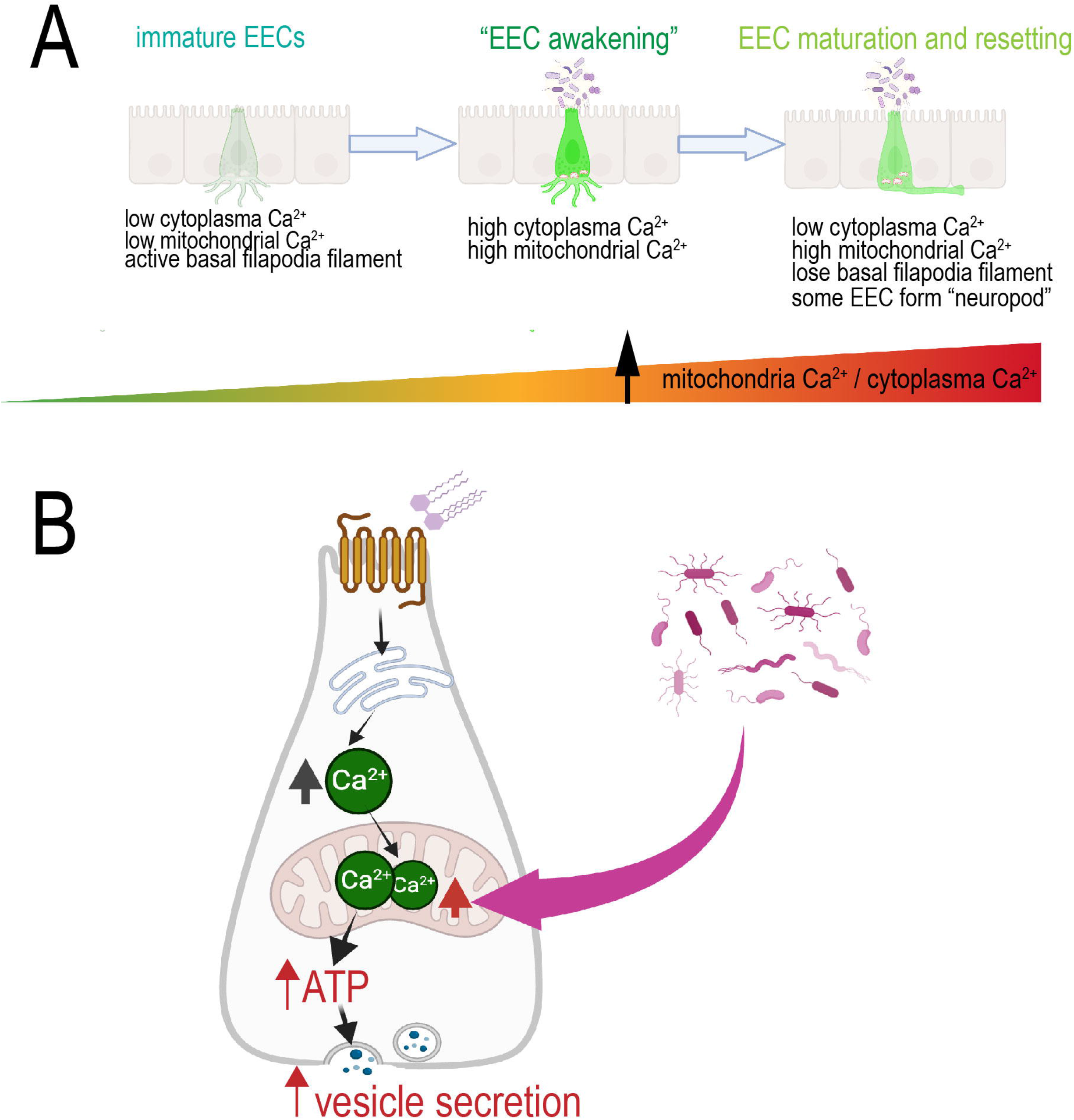
Model figure showing gut microbiota modulate EEC maturation and mitochondrial function. (A) During early development, the immature EECs exhibit low cytoplasmic Ca^2+^ and low mitochondrial Ca^2+^ levels. These immature EECs have active filapodial filaments at the basal lateral membrane. After the zebrafish hatched out and commensal microbiota started to colonize the zebrafish intestine, the EECs continued to develop and mature. Shortly after commensal microbiota colonization, the EECs increase both cytoplasmic and mitochondrial Ca^2+^ significantly (“EEC awakening”). After the EEC awakening, the EECs continue to mature and lose the basal lateral filapodial filaments. Some EECs form a neuropod. The mature EECs have low cytoplasmic Ca^2+^ but high mitochondria-to-cytoplasm Ca^2+^ ratio. (B) Commensal microbiota promotes EEC mitochondrial respiration function and increases mitochondrial inner membrane electronic potential (ΔΨm). When nutrient stimulants, like fatty acids, stimulate the EECs, the EEC cytoplasmic Ca^2+^ rises. The high ΔΨm permits the cytoplasmic Ca^2+^ to flux into the mitochondrial matrix and power mitochondrial ATP production, which then promotes EEC vesicle release.

